# Cytoskeletal organization in isolated plant cells under geometry control

**DOI:** 10.1101/784595

**Authors:** P. Durand-Smet, Tamsin A. Spelman, E. M. Meyerowitz, H. Jönsson

## Abstract

Specific cell and tissue form is essential to support many biological functions of living organisms. During development, the creation of different shapes at the cellular and tissue level fundamentally requires the integration of genetic, biochemical and physical inputs.

It is well established that the cortical microtubule network plays a key role in the morphogenesis of the plant cell wall by guiding the organisation of new cell wall material. Moreover, it has been suggested that light or mechanical stresses can orient the microtubules thereby controlling wall architecture and plant cell shape. The cytoskeleton is thus a major determinant of plant cell shape. What is less clear is how cell shape in turn influences cytoskeletal organization.

Recent *in vitro* experiments and numerical simulations predicted that a geometry-based rule is sufficient to explain some of the microtubule organization observed in cells. Due to their high flexural rigidity and persistence length of the order of a few millimeters, MTs are rigid over cellular dimensions and are thus expected to align along their long axis if constrained in specific geometries. This hypothesis remains to be tested *in cellulo*.

Here we present an experimental approach to explore the relative contribution of geometry to the final organization of actin and microtubule cytoskeletons in single plant cells. We show that, in cells constrained in rectangular shapes, the cytoskeleton align along the long axis of the cells. By studying actin and microtubules in cells with the same system we show that while actin organisation requires microtubules to be present to align the converse is not the case. A model of self organizing microtubules in 3D predicts that severing of microtubules is an important parameter controlling the anisotropy of the microtubule network. We experimentally confirmed the model predictions by analysing the response to shape change in plant cells with altered microtubule severing dynamics. This work is a first step towards assessing quantitatively how cell geometry contributes to the control of cytoskeletal organization in living plant cells.

## Introduction

How organisms achieve their specific forms and alter their growth patterns in response to environmental signals has been the subject of investigation for over a century (Thompson 1917). The importance of shape for specific biological functions has been shown across kingdoms, for example: the shape of some bacteria facilitates motility (Mitchell 2002) or allows them to avoid phagocytosis (Champion and Mitragotri 2006). The shape of diatoms can facilitates nutrient uptake (Pahlow et al. 1997). The aperture and closure mechanism of pores allowing gas exchange between plants and the environment relies on the shape of stomatal guard cells in leaves of flowering plants (Galatis and Apostolakos 2004; Franks and Farquhar 2007). Or again, the elongation of cells promotes a pro-healing phenotype in macrophages (McWhorter et al. 2013).

The cytoskeleton, an interconnected network of filamentous polymers such as actin and microtubules (MTs), plays a key role in establishing robust cell shape. In animal cells, the actomyosin cortex gives the cells their mechanical strength and acts in controlling their shapes (Fabry et al. 2003; Balland et al. 2005; Smith et al. 2005; Fletcher and Mullins 2010). In prokaryotes, the actin homologous protein MreB is involved in determining cell shape by guiding cell wall synthesis (Werner et al. 2009; L. J. Jones et al. 2001; Figge et al. 2004). In plant cells, the cortical MT network prescribes the shape of cells by guiding the synthesis of cellulose in the cell wall (Paredez et al. 2006) and supports the mechanical strength of plant protoplasts (plant cells without walls) (Durand-Smet et al. 2014). In growing plant cells, the actin network is also involved in cell wall formation by being responsible for global distribution of cellulose synthase complexes at the plasma membrane (Sampathkumar et al. 2013).

Many external signals such as light or hormones can influence MT direction (Zandomeni and Schopfer 1993). In addition, many studies suggest that mechanical stresses, generated by tissue shape and turgor pressure as well as by external application of force, orient the MTs along their principal direction in the shoot apical meristem (SAM) (Hamant et al. 2008), in leaf cells (Sampathkumar, Yan, et al. 2014; Jacques et al. 2013) in hypocotyls (Verger et al. 2018; Robinson and Kuhlemeier 2018; Hejnowicz et al. 2000), in sepals (Hervieux et al. 2017; Hervieux et al. 2016) or in single protoplasts (Wymer et al. 1996). The response of MTs to mechanical stress relies on dynamic self-organization of the MTs. In particular, MT rearrangement after changes in mechanical stress is promoted by the microtubule-severing protein katanin (Uyttewaal et al. 2012; Sampathkumar, Krupinski, et al. 2014). The reorganization of MTs after blue light exposure or auxin treatment also depends on severing by katanin (Chen et al. 2014; Lindeboom et al. 2013). As cells grow, changes in mechanical stress are hypothesized to influence cytoskeletal organization, which further influences wall anisotropy and local growth rates, creating a feedback loop that could be exploited to generate specific cell shape and tissue form. The cell cytoskeleton is often viewed as a major determinant of cell shape. While the impact of geometry on cytoskeletal organization is well characterized in animal cells (Gomez et al. 2016),(Théry et al. 2006; Versaevel et al. 2012); (Bao et al. 2017; Bao et al. 2019) and in *in vitro* systems (Cosentino Lagomarsino et al. 2007; Soares e Silva et al. 2011; Alvarado et al. 2014; Pinot et al. 2009), single cell analysis *in planta* is lacking. Since plant cells are encased into a stiff wall, strong and persistent geometrical constraints act on cells throughout the plant life, the importance of cell geometry for organising cytoskeletal filaments in living plant cells needs to be better understood.

*In planta*, different MT organizations can be observed in cells with similar geometry within the same tissue, for example in epidermal regions within the shoot apical meristem (SAM) or in different cell layers in the hypocotyl of *Arabidopsis thaliana* plants. Because shape-derived mechanical stresses play a dominant role in this case, it is hard to understand the role of cell geometry alone on cytoskeletal organization when working at the tissue level. To overcome this issue, recent studies used 3D modelling to explore the impact of cell geometry on MT organization in plant cells (Mirabet et al. 2018; Chakrabortty, Blilou, et al. 2018; Chakrabortty, Willemsen, et al. 2018). Basic MT-MT interaction rules coupling MT dynamics to the geometry of the cell were shown to be sufficient to explain the division patterns observed experimentally in embryonic cells (Chakrabortty, Willemsen, et al. 2018). In this study, the direction of maximal wall stress in orienting the MTs was not required. The simulations from Mirabet et al. showed that the preferred longitudinal organization of MTs in elongated shapes spontaneously emerges from the dynamical properties of the MT array and the bending rigidity of the MT rod filaments (Mirabet et al. 2018). This geometry-driven organization was overcome by introducing a small directional bias, suggesting that the geometry-driven rule could be weak in a context of growing plant cells, allowing for outside mechanical forces to alter cytoskeletal organization.

These studies suggest that geometry could be a sufficient rule for MT organization; nevertheless this hypothesis needs to be addressed experimentally in the context of real living cells. Moreover, severing of MTs by katanin proteins has not been incorporated into previous fully 3D numerical simulations. The MT organization emerging from simulations when incorporating the severing rule, and whether that can still summarize experimental observations, is not clear.

In the context of studying the MT organization in response to geometrical confinement in plant cells, it is of interest to correlate this with the response of the actin network. More specifically, it is well known that actin organization is influenced by mechanical forces in animal cells (Harris et al. 2018; Balaban et al. 2001; Wang et al. 2001), but how actin filaments respond to mechanical stresses has rarely been investigated in plants (Goodbody and Lloyd 1990; Hardham et al. 2008; Branco et al. 2017). Because the actin filaments and MTs have been shown to localize near the plasma membrane in plant cells it is believed that the two networks interact (Traas 1984). The specific interactions between actin and MTs are poorly understood, but one study demonstrated that actin filament reassembly depends on MTs (Sampathkumar et al. 2011). This interaction might be facilitated by plant formins, proteins capable of binding both actin filaments and MTs (Sampathkumar et al. 2011; Wang et al. 2012; Deeks et al. 2010). Whether actin filaments and MTs interact in the context of a response to geometrical or mechanical perturbation still remains to be established.

In this study, we developed an experimental approach to explore the contribution of geometry to the final organization of actin and microtubule cytoskeletons in isolated plant cells without cell walls. We extended a previously developed model of 3D self-organizing MTs network to predict the role of severing of microtubules in the shape response. We show that geometry alone is sufficient to explain the observed reorganization of MTs in rectangular shapes, and that the geometry response is dependent on severing. In addition, studying actin and microtubules in the same system allowed us to compare the organization of the two networks, and how they interact. We show that the actin response to shape change depends on the MT network.

## Results

To study how geometry affects cytoskeletal organization in plant cells we developed an experimental approach allowing the control of the geometry of cells. The approach has been successfully used in the past to control the geometry of sea urchin embryos for example (Minc et al. 2011). The principle consists of confining single cells into microfabricated microwells of various geometries.

Because we aim to analyse the effect of geometry independently from mechanical stresses in the wall and from tissue signals, plant cell walls were digested and single protoplasts were used for experiments. In order to be properly fitted into the micro-wells, the osmotic pressure of the protoplasts was reduced to a level which would correspond to plasmolysis in a context of cell with a wall (fig1, Methods). We used green-fluorescent protein fused to a microtubule binding domain (MBD-GFP) (Granger and Cyr 2001), (Marc et al. 1998) and green fluorescent protein fused to an F-actin binding domain (FABD-GFP) (Ketelaar et al. 2004) reporter lines for monitoring the microtubules and the actin network, respectively. The microwells were cast from PCM medium with agarose (Methods) with wells of various shapes and a set of diameters ranging from 15 to 40µm.

**Figure 1:**
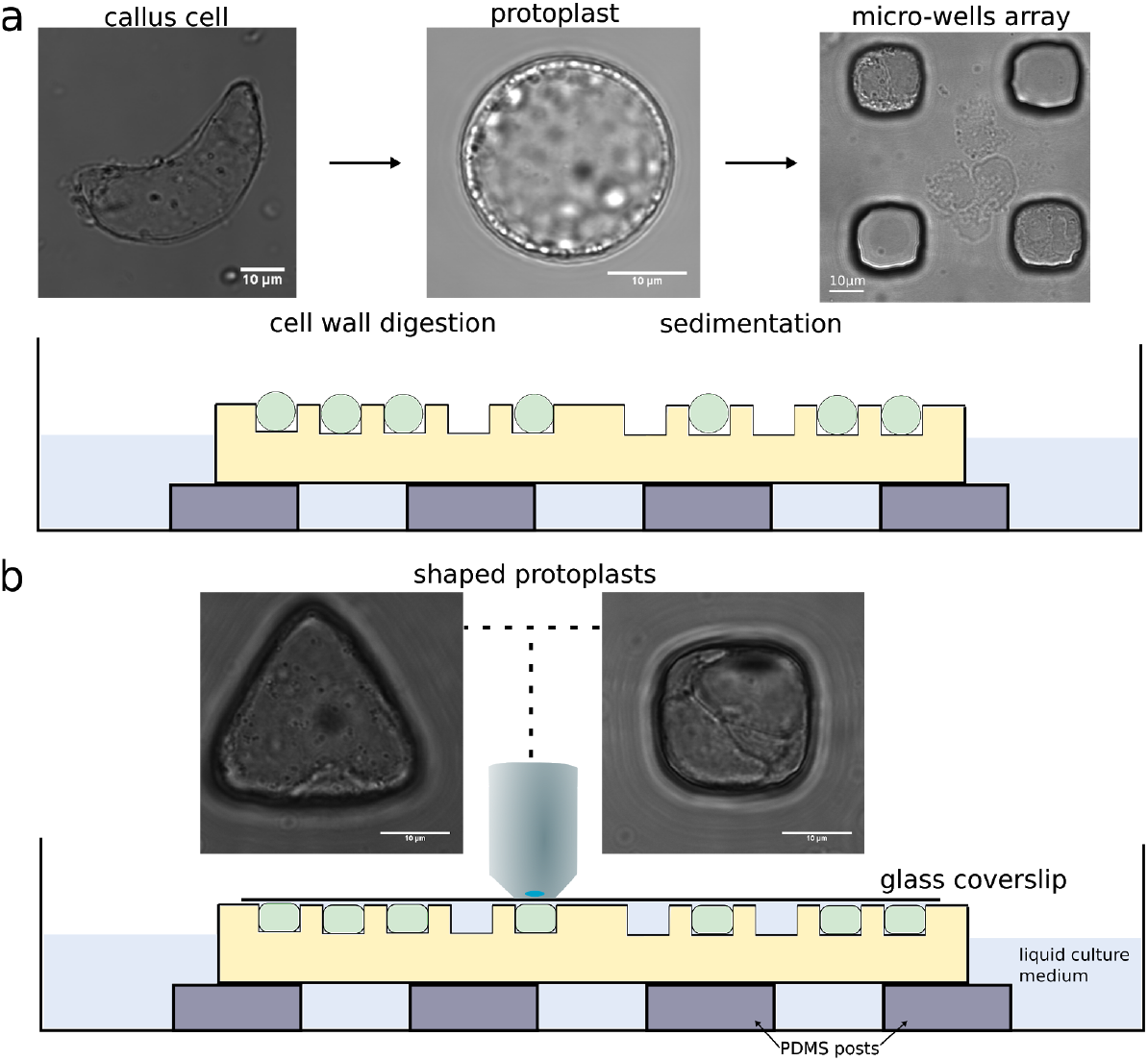
Description of the experimental set-up used for the confinement of protoplasts in different geometries. (a) The cell walls of Arabidopsis thaliana root-derived callus cells were digested to make spherical protoplasts. Protoplasts were then plated on top of the microwell array. Left: bright field picture of an isolated cell from callus with undefined geometry, middle: bright field picture of a freshly isolated protoplast with a spherical geometry, right: bright field picture of protoplasts in square micro-wells after sedimentation. Bottom: illustration of microfluidic setup with cells loaded in wells. (b) Coverslips were carefully placed on top of the microwells and the samples were imaged using an upright microscope. Bright field pictures of protoplasts being shaped in microwells of various geometries (x63 objective), All size standards are 10µm.

### Actin organization in protoplasts confined in controlled geometries

We first tested the effect of geometry on the organization of cytoskeletal networks in unperturbed protoplasts. For that purpose, we chose to focus on microwells with simple shapes with in-plane geometry of circles, squares, triangles or rectangles (figS1). Cells were observed between 30min and 5h after being fitted in the microwells. The average angle and the anisotropy of the actin (fig2-a) and MT (fig2-d) networks in protoplasts in those shapes were then quantified (Methods).

**Figure 2:**
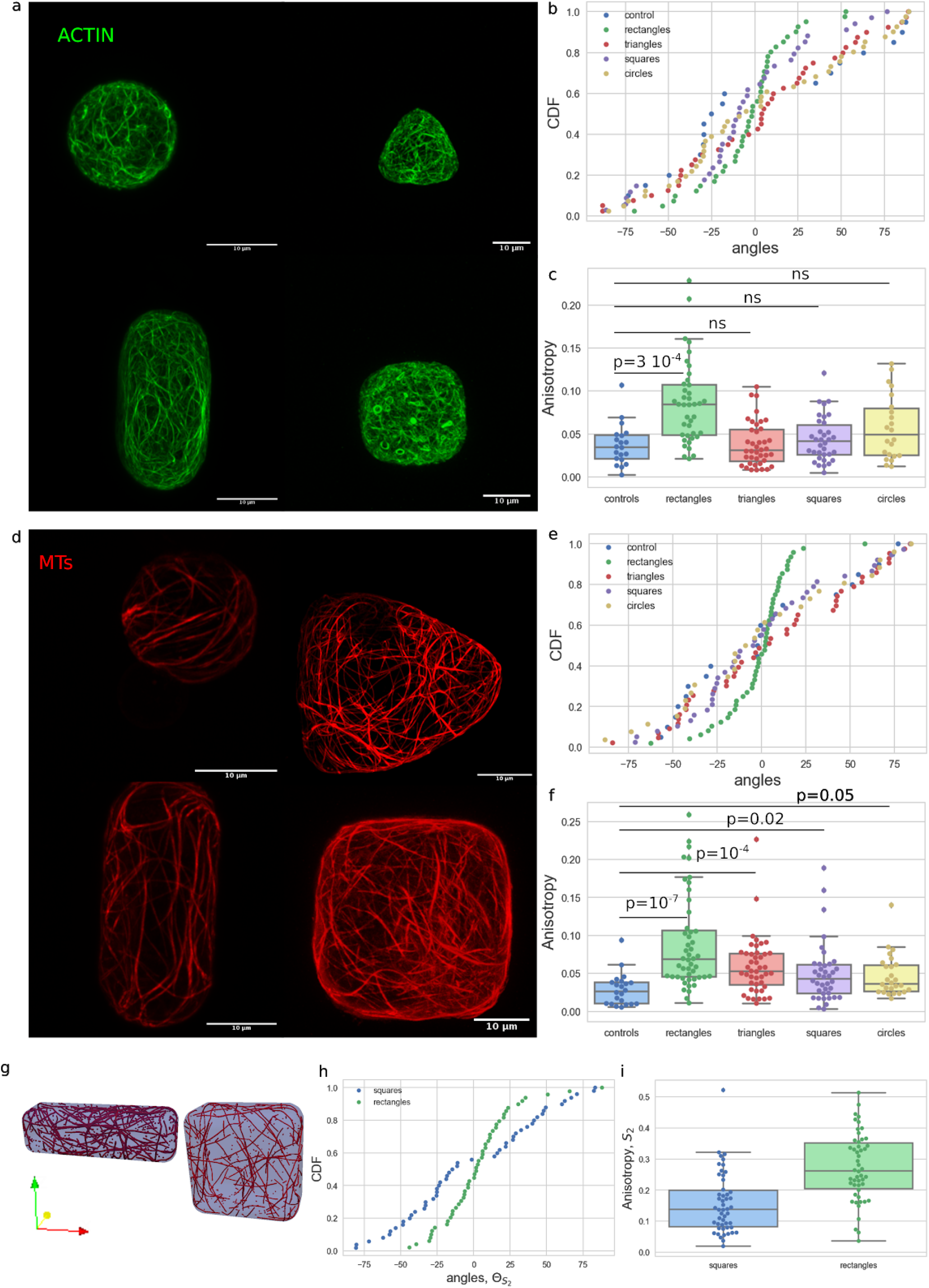
Effect of the shape on the actin and microtubule network organization. (a) Actin organization: protoplasts expressing FABD-GFP reporter confined in microwells of different geometries showing the effect of shape on the actin network organization in protoplasts. (b) cumulative distribution of the average angle of the actin network for the various geometries tested (see also figS2). Number of protoplasts N=40 and number of repeat n=11 for triangles, N=40 and n=12 for rectangles, N=34 and n=12 for squares, N=22 and n=3 for circles, N=40 and n=2 for controls. (c) boxplot of the anisotropy of the actin network in the different geometries tested. (d) MT organization: protoplasts expressing MBD-GFP reporter confined in microwells of different geometries showing the effect of shape on the MT network organization in protoplasts. (e) Cumulative distribution of the average angle of the MT network for the various geometries tested (see also figS2 and S3). Number of protoplasts N=44 and number of repeat n=12 for triangles, N=48 and n=10 for rectangles, N=38 and n=11 for squares, N=26 and n=7 for circles, N=22 and n=4 for controls. (f) Boxplot of the anisotropy of the MT network in the different geometries tested. (g) example output of the numerical simulations of MTs for a square domain and a rectangle domain. (h) graph of the average angle of MT network and (i) anisotropy found for rectangular and square shapes in numerical simulations for 10,000 time steps and *P*_*cross*_ = 0.005 (with severing of microtubules). All size standards are 10µm

Our experiments suggest that the average orientation of the actin network is not influenced by geometry for cells in circular, triangular and square shapes (fig2b, figS2). The distributions of the average angle showed no preferred actin orientation in cells in these shapes and were comparable in uniformity to the distribution found in spherical protoplasts (the control shape; p-values>0.05 with Kupier test, Methods, FigS2). The anisotropy (measured as the difference of the eigenvalues of the nematic tensor calculated from the intensity gradients in individual cells, Methods) of the actin network in triangular, square and circular shapes is not significantly different than the anisotropy of the actin network in spherical protoplasts (fig2c, KS test gave p-values>0.05).

In contrast, the average orientation of the actin network was found to be strongly biased toward the long axis in protoplasts in rectangular shapes (fig2b). The cumulative distribution of the average angle for the actin network in protoplasts confined in rectangular shapes shows an accumulation around the 0º degree angle, which corresponds to the long axis of the rectangle (fig2b and S2, p-value Kuiper=8 10^−6^). Moreover, the anisotropy of the actin network in rectangular shapes is significantly higher than for the control shape, suggesting that the actin filaments are better aligned to one another in the rectangular shapes (p-value KS = 3 10^−4^). In the conditions of our experiments, actin follows cell shape as in vitro (Soares e Silva et al. 2011; Alvarado et al. 2014) indicating that intrinsic mechanical properties of actin filaments dictates their orientation.

### MTs organization in protoplasts confined in controlled geometries

We next quantitatively analyzed MT organization. We found that the average orientation of the MT network is not biased in circular, triangular and square shapes (fig2e and S2 and S3). Similarly to actin, the distributions of the average angle showed no preferred orientation over all of the cells tested in these shapes and the distributions are the same as that for the spherical protoplasts. However, the anisotropy of the MT network in cells of those shapes is always significantly higher than the anisotropy of the MT network in spherical protoplasts (fig2f KS p-values of 10^−7,^ 10^−4^ 0.02 or 0.05 for rectangular, triangular, squares and circular shapes respectively). This suggests that the confinement alone is enough to influence the alignment of MTs. This was also observed *in vitro* when microtubules were allowed to organize freely in droplets of different sizes (Pinot et al. 2009).

Similarly to what was found for actin, the average orientation of the MT network was strongly biased towards the long axis in protoplasts in rectangular shapes (fig2e and S2). The cumulative distribution of the average angle for the MT network in protoplasts confined in elongated shapes shows an accumulation around the 0 degree angle (which corresponds to the long axis of the shape)(fig2e and S2, p-value Kuiper = 10^−16^). The anisotropy of the MT network in elongated shapes is also significantly higher than for the control, suggesting that the MTs are better aligned to one another in the elongated shapes (fig2f, p-value KS test=10^−7^). These results in living single plant cells provide experimental evidence to the specific MT organization that was predicted *in silico* via numerical simulations (Mirabet et al. 2018) and *in vitro* (Cosentino Lagomarsino et al. 2007; Cortès et al. 2006).

In most numerical simulations performed on self organizing MTs, the dynamical properties of the MTs were restricted to a small number of parameters and some aspects of MT growth were not considered, such as severing of microtubules by katanin in the 3D simulations (Mirabet et al. 2018). Recently, an extensive study of the effect of severing of MTs highlighted the importance of taking into account this dynamical property in simulations (Deinum et al. 2017). However in this study, simulations were restricted to surfaces and did not account for the full 3D nature of the MT network. To study this, we extended a previously developed 3D model (Mirabet et. al. 2018) by adding crossover severing to the MT interactions (Methods, figS4). Briefly, we modeled each MT as a line segment with the segment growing, shrinking and interacting with other segments. We imposed the interaction rules that when MTs meet at an angle below 40° they bundle; for angles over 40° there is an induced catastrophe (at each tubulin subunit) with probability P_cat_. If no induced catastrophe occurs the MT keeps growing but will be severed with a probability P_cross_ at any later step if both the crossing and crossed-over tubulin still exist (figS4). The 3D model with severing leads to alignment of the MT network with the long axis of rectangular shapes (fig2g-h). Contrary to what was found before in square domains without severing of the MTs (Mirabet et al. 2018), the simulations with our severing rules led to no significant preference to align along the diagonals of the square. The experiments also show no significant alignment along the diagonals. Together, these results illuminate the role played by severing in self-organizing MTs networks. In addition, the results of the simulations show that the anisotropy of the network is higher in rectangular domains (fig2i) than in square domains which is in agreement with our experiments. Because similar organization of the MT network in elongated shapes was found in numerical simulations based on intrinsic dynamical properties of MTs, our results suggest that geometry might be a sufficient rule to organize the MTs in some living plant cells.

Next, to better compare the actin and MT networks organizations, we did an independent set of experiments with higher imaging resolution, focussing on rectangular shapes (and their control, spherical protoplasts) as those produced the greatest effect (figS5). These set of experiments resulted in an overall increase in anisotropy values due to reduced signal to noise ratio yet showed a similar trend as the previous experiments. The distributions for the average angles in all of the cells tested show that the actin and MT average angle is distributed around the 0º angle which corresponds to the long axis of the rectangular shapes (p-value = 5 10^−7^ for the actin and 4 10^−4^ for the MTs, figS5). In contrast, the average angle distribution does not show any specific bias for the spherical protoplasts and the distributions are comparable to uniform distributions (p-value = 0.6 for the actin and 0.6 for the MTs, figS5). The anisotropy is always higher for the MT network than for the actin network (figS5), which may reflect different rigidity of MT and actin filaments (Gittes et al. 1993).

### Actin organization in protoplasts confined in elongated geometry relies on the microtubule network but not vice versa

Since our results suggest that actin filaments and MTs organize similarly in rectangular shapes we next asked if the actin and MTs networks interact with each other in isolated plant protoplasts under geometrical constraints.

To investigate this, we first studied the organization of the actin network when the MTs were depolymerized with oryzalin (fig3a and figS6). In this condition, the average angle of the actin network in protoplasts in elongated shape was still aligned with the long axis of the shape, meaning that absence of MTs had no significant effect on the orientation of actin after cell shape change (fig3b and figS5). In contrast to the untreated case, for protoplasts treated with oryzalin, the anisotropy of the actin network was not significantly higher in rectangular shapes than in control (fig3c, cf. fig2c). This indicates that the MT alignment is required for the actin filaments to align well to one another in response to shape.

**Figure 3:**
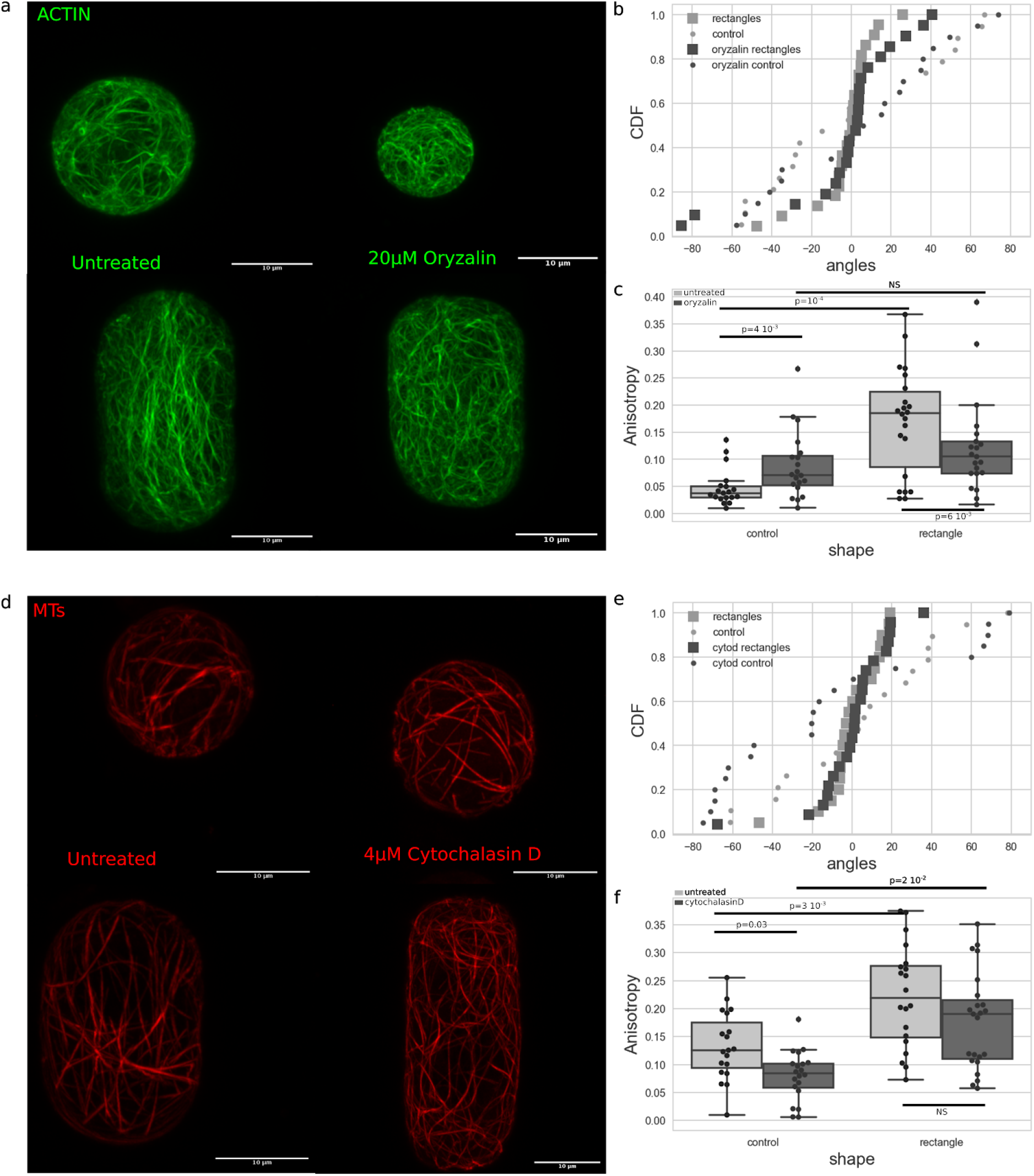
(a) Actin organization: protoplasts expressing FABD-GFP reporter confined in microwells showing the effect of shape on the actin network organization in untreated protoplasts (left) and protoplasts treated with Oryzalin (right). (b) Cumulative distributions of the average angle of the actin networks of untreated protoplasts (light grey) and protoplasts treated with 20μm oryzalin (dark grey) in rectangular microwells (square markers) and in control-shaped (circle markers) (see also FigS7). Number of protoplasts N=19 and number of repeat n=1 for untreated protoplasts in control shapes, N=22 and n=4 for untreated protoplasts in rectangular shapes, N=20 and n=2 for treated protoplasts in control shapes, N=21 and n=2 for treated protoplasts in rectangular shapes. (c) Boxplot comparing the anisotropy of the actin networks of untreated protoplasts (light grey) and protoplasts treated with 20μm of oryzalin (dark grey). (d) MT organization: protoplasts expressing MBD-GFP reporter confined in microwells showing the effect of shape on the MT network organization in untreated protoplasts (left) and protoplasts treated with CytochalasinD (right). (e) Cumulative distributions of the average angle of the MT networks of untreated protoplasts (light grey) and protoplasts treated with 4μm cytochalasin D (dark grey) in rectangular microwells (square markers) and in control-shape (circle markers) (see also figS7). Number of protoplasts N=19 and number of repeat n=3 for untreated protoplasts in control shapes, N=20 and n=3 for untreated protoplasts in rectangular shapes, N=20 and n=2 for treated protoplasts in control shapes, N=23 and n=1 for treated protoplasts in rectangular shapes. (f) Boxplot comparing the anisotropy of the MT networks of untreated protoplasts (light grey) and protoplasts treated with 4μm cytochalasin D (dark grey). All size standards are 10µm

We then studied the organization of the MT network when actin was depolymerized with cytochalasin D (protoplasts expressing the MBD-GFP reporter were treated for 1h with 4μM of CytoD) (fig3d and figS6). In this condition, the average angle of the MT network in protoplasts in elongated shape was aligned with the long axis of the shape, meaning that depolymerization of the actin has no effect on the reorganization of MTs upon shape change in terms of preferred angle (fig3e and figS5). The anisotropy of the MT network still increased in protoplasts plated in elongated shape as compared to spherical protoplasts (fig3f, p-value = 0.02). Thus, the disruption of the actin network did not have a significant impact on the response of MTs to shape in the conditions of our experiments.

Our findings suggest that the actin organization relies on the pre-existing MT organization but not vice versa.

### Actin polymerization defect impairs the organization of actin in response to shape

We next try to challenge the robustness of the actin response to shape by testing if an alteration in actin polymerization affected its organization in response to shape. The *act2* mutant is known to show disrupted actin polymerization and cells in the mutant exhibit shorter actin filaments (Nishimura et al. 2003; Vaškebová et al. 2018). We performed experiments on protoplasts of *act2-5* mutants expressing the FABD-GFP reporter for actin (fig4a-b). The average angle for the actin network was found to be aligned to the main axis of the shape for a majority of *act2-5* protoplasts confined in rectangular shape and the average angle was found to be uniformly distributed for spherical *act2-5* protoplasts. This is similar to what was found for the wild type protoplasts. It seems that this defect in polymerization does not affect the ability of the actin network to properly orient in response to shape. Nevertheless, when looking at the anisotropy of the actin network we found that *act2-5* protoplasts failed to respond to rectangular shape. In fact, the distribution of the anisotropy of the actin network for *act2-5* protoplasts in rectangular shape is not significantly different compared to the distribution in spherical *act2-5* protoplasts (p-value=0.2). Moreover, we found that actin in *act2-5* spherical protoplasts exhibited higher anisotropy than in the wild type spherical protoplasts. Higher anisotropy is thus a consequence of the defect in polymerization in the *act2-5* mutants. Altogether, our results suggest that while the long axis is selected in the mutant, actin polymerization and filament length are important for the magnitude of the cell shape response.

**Figure 4:**
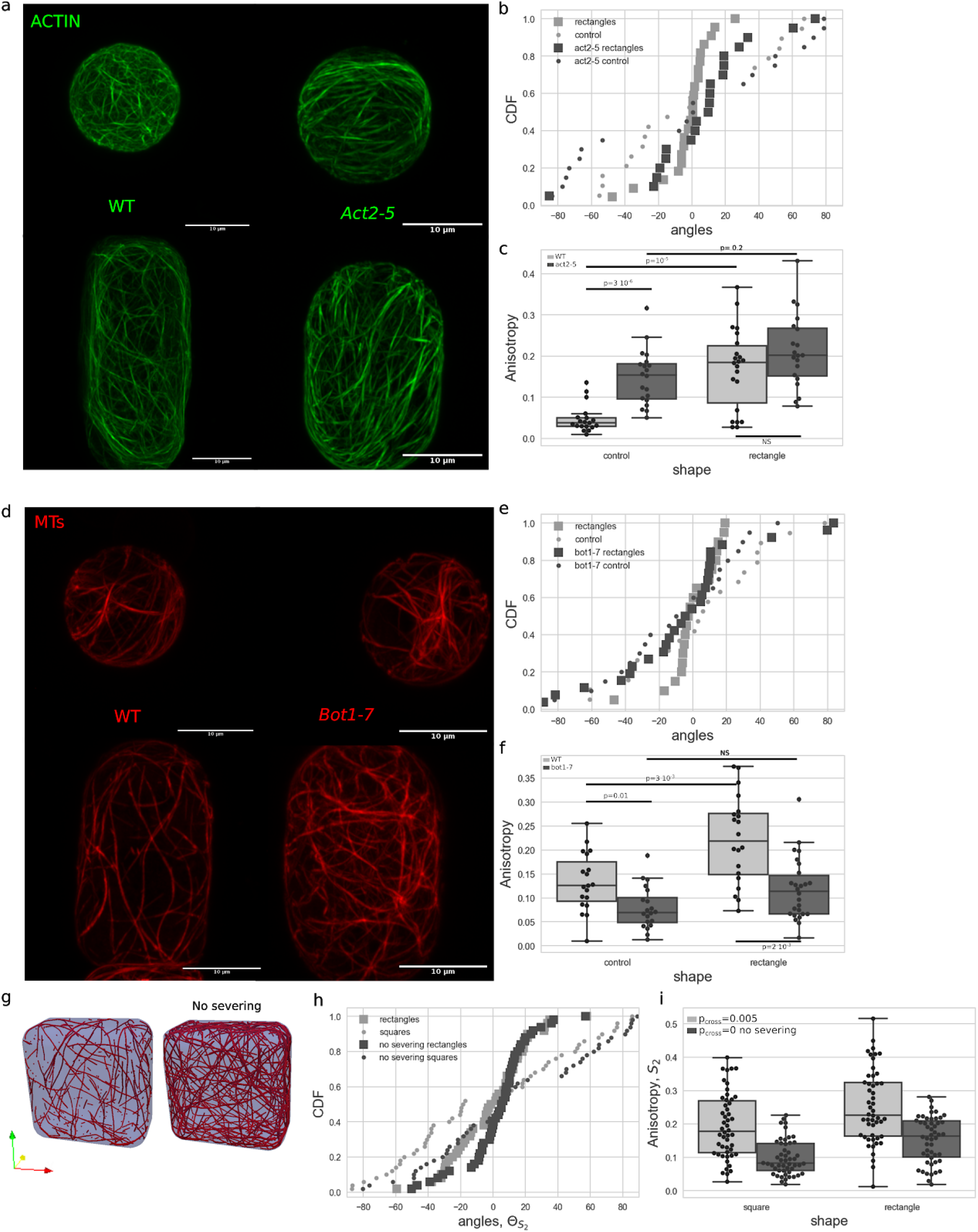
shape response in protoplasts with genetically disturbed cytoskeleton dynamic. (a) Actin organization: protoplasts expressing FABD-GFP reporter confined in microwells showing the effect of shape on the actin network organization in wild type protoplasts (left) and protoplasts generated from callus in the *Act2-5* mutation background (right). (b) Cumulative distributions of the average angle of the actin networks of wild type protoplasts (light grey) and protoplasts generated from callus in the *Act2-5* mutation background (dark grey) in rectangular microwells (square markers) and in control shape (circle markers) (see also figS8). Number of protoplasts N=19 and number of repeat n=1 for wild type protoplasts in control shapes, N=22 and n=4 for wild type protoplasts in rectangular shapes, N=20 and n=2 for protoplasts in the *Act2-5* background in control shapes, N=20 and n=2 for protoplasts in the *Act2-5* background in rectangular shapes. (c) Boxplot comparing the anisotropy of the actin networks of wild type protoplasts (light grey) and protoplasts generated from callus in the *Act2-5* mutation background (dark grey). (d) MT organization: protoplasts expressing MBD-GFP reporter confined in microwells showing the effect of shape on the MT network organization in wild type protoplasts (left) and protoplasts generated from callus in the *bot1-7* mutation background (right). (e) Cumulative distributions of the average angle of the MT network of wild type protoplasts (light grey) and protoplasts generated from callus in the *bot1-7* mutation background (dark grey) in rectangular micro-wells (square markers) and in control-shaped (circle markers) (figS8). Number of protoplasts N=19 and number of repeat n=3 for wild type protoplasts in control shapes, N=20 and n=3 for wild type protoplasts in rectangular shapes, N=20 and n=1 for protoplasts the *bot1-7* background in control shapes, N=26 and n=1 for protoplasts in the *bot1-7* background in rectangular shapes. (f) Boxplot comparing the anisotropy of the MT networks of wild type protoplasts (light grey) and protoplasts generated from callus in the *bot1-7* mutation background (dark grey). (g) Example time point from numerical simulation of the MTs with (left) and without crossover severing (right). (h) graph of the average angle of MT network and (i) anisotropy found for rectangular and square shapes in numerical simulations with (P_cross_=0.005) and without (P_cross_=0) crossover severing for each case averaged over 50 simulations after 3000 timesteps. All size standards are 10µm

### Severing of microtubules by katanin is necessary for the MT response to shape

We next tested whether an alteration in MT dynamics affected MT organization in response to shape. Several studies suggest that plant MT self-organization depends on katanin (Uyttewaal et al. 2012). This protein is known to promote severing of MTs at the sites of MT encounters and thus promotes the alignment between MTs (Lindeboom et al. 2013) (Deinum et al. 2017).

To test whether katanin activity is required for this MT response, we performed experiments on protoplasts expressing the MBD-GFP reporter in the katanin mutant background *bot1-7* (fig4d). We found that the average angle of the MT network was still distributed following the long axis of the shape but that the number of cells with a transverse organization was increased in comparison to wild type. For the wild type, almost all cells exhibited an average angle between −20 and 20 degrees (fig4e, n=19/20) whereas in the mutant more than one third of the cells exhibited an average angle higher than 20 degrees (n=10/26) and some cells had a dominating transverse array of MTs (n=5/26). Moreover, the distribution of the average angle for the MTs in the mutant protoplasts in rectangular shapes was no longer different compared to a uniform distribution (fig4e and S8, Kuiper test gives a p-value higher than 0.05).

In addition, in the *bot1-7* mutant, the anisotropy of the MT network was smaller than in wild type in spherical protoplasts (fig4f). This defect in anisotropy of the network for this mutant is already known from *in vivo* studies (Burk et al. 2001; Bichet et al. 2001). In protoplasts from *bot1-7* mutant cells, the anisotropy of the MT network was not significantly higher in the protoplasts plated in rectangular wells when compared to spherical protoplasts (fig4f). While the shape creates a bias toward a better alignment of MTs in the wild type, this bias no longer exists in the absence of the action of katanin. Our experiments suggest that a katanin mutant is less responsive to geometrical cues.

In the simulations we tested different values of the parameter representing the severing by katanin (fig4g-h). Our computational simulations demonstrated that increasing the probability that crossing MTs will be severed, *P*_*cross*_, increases the alignment of the MTs (fig4i, figS9). Since katanin is a MT-severing protein, we expect higher concentrations of katanin to correspond to larger *P*_*cross*_. Indeed, the computational results demonstrate that higher *P*_*cross*_can cause more anisotropy which corroborates our experimental findings that katanin is required for MT organisation.

## Conclusion/Discussion

In this study, we quantified cytoskeletal response to cell shape using a novel approach. By working with isolated protoplasts in reduced osmotic conditions we diminished possible tissue signals including the stress and strain patterns resulting from tissue shape and turgor pressure, and eliminated the effects of tension usually borne by the cell wall. In these conditions we expect to be as close as possible to study the effect of geometry alone. We show that MTs and actin filaments are sensitive to cell geometry in living isolated plant cells and that they align mostly along the long axis of cells. A model based on basic MT interaction rules and on intrinsic mechanical properties of MTs is sufficient to explain our experimental quantification. Therefore, we conclude that, at the single cell level, geometry is a contributor to cytoskeletal organization in the plant cells we tested.

We found that both actin and MT networks exhibited a specific organization only when the cells were confined in rectangular wells. Nevertheless, for triangular and square shapes, while we did not see a significant bias of the shape on the organization, we found that for a small proportion of cells (5/30, figS3) MTs were aligned along the edges of triangular shapes or presented a weak bias toward the diagonals of square shapes (13/38 cells, figS3). Contrary to what was found in animal cells for the actin (Bao et al. 2017; Théry et al. 2006) or predicted with numerical simulations of self-organizing MTs (Mirabet et al. 2018), we find only a weak bias toward diagonals in our experiments and simulations. One explanation might be that the square shapes, due to technical limitations in the experiments, do not have sharp corners. Square wells are not drastically distinct from circular wells, thus the shape might not be distinct enough to induce a strong enough bias.

In rectangular shapes, both actin and MTs were found to be aligned along the long axis and the anisotropy of the networks was always higher than in spherical protoplasts. This is similar to what was found *in vitro* (Soares e Silva et al. 2011; Alvarado et al. 2014; Cosentino Lagomarsino et al. 2007; Cortès et al. 2006) suggesting that the organization emerges from intrinsic mechanical properties of the network, a self-organizing process. Comparing actin and MTs organization in the same system revealed that the anisotropy of the actin network is always smaller than for the MTs. This might be linked to the different nature of the filaments composing each network (MT being an association of dimers organized in tubes and actin being a linear composition of monomers). *In vitro*, while the MTs are rather stiff rods, the flexural rigidity of actin filaments is 1000 times smaller than for MTs (Gittes et al. 1993). That could explain why the actin filaments can adopt more configurations in the same amount of space available and thus the alignment is not as robust as for the MTs.

*In vivo*, transverse organization of MTs is quite often observed in elongated plant cells, an orientation we did not observe in our system in wild type protoplasts in rectangular shapes. It has been shown in numerical simulations that a small bias in the growth direction of MTs is sufficient to alter this longitudinal organization into a transverse one (Mirabet et al. 2018). Similarly, while both transverse and longitudinal populations can be found in numerical simulations, the probability of catastrophe at the edges of the domain can create a bias toward one orientation (Chakrabortty et al. 2018). This suggests that the geometry of the cells has only a relative influence on cytoskeletal organization *in planta* and that in the absence of bias the steric constraint due to the mechanical properties of MTs prevails.

A recent perspective paper proposed that MTs possibly can directly sense tension accumulating in the cell wall and align accordingly (Hamant et al. 2019). The alignment of MTs with the maximum tensile stress direction would create a favored stable configuration for MTs growth. Re-orientation of MTs to the longitudinal axis in leaf epidermal cells concomitant with the detachment of MTs from the plasma membrane was previously observed (Sainsbury et al. 2008). The mechanical tension accumulating in the cell wall can be transmitted to the MTs only if the wall-membrane-MT continuum is intact. Because there is no cell wall in our case and because we worked with protoplasts in reduced osmotic conditions, we hypothesize that the default geometry rule dominates the response in our experimental system. Nevertheless, it is noteworthy that constraining a spherical cell into a different geometry creates mechanical stresses in the plasma membrane. Because the plasma membrane is believed to behave mostly like a fluid material, it is not clear whether these mechanical stresses are fully relaxed in our experiments or if that tension is transmitted to the cytoskeletal networks.

We did not study the dynamics of cytoskeletal reorganization process from spherical shape to rectangular shape. *In vitro*, an external geometry can strongly influence spatial organization of semiflexible filaments if the dimension of the confinement is comparable to the persistence length of the filaments. For example, actin filaments align parallel to the long walls of microchambers as soon as the geometrical aspect ratio of the chamber is greater than 1.5, which indicates that the organization of the filaments is dominated by geometrical boundary interactions (Alvarado et al. 2014). We briefly investigated this with the available quantification of our experimental data and we found that there is no apparent correlation between the aspect ratio of the cell and average orientation of the network in cells confined in rectangles (figS10).

Our study provides evidence that actin filaments and MTs interact in the context of a response to geometrical perturbation. We find that actin organizes longitudinally in elongated shapes independently of MTs but that the level of alignment between actin filaments in those shapes depends on the MT cytoskeleton. It was already reported that actin distribution becomes more irregular when the MTs are depolymerized (Traas et al. 1987). In our experiments, depolymerizing actin did not reveal any disturbed MT organization in cells of elongated shapes, suggesting that MT organization is in this case independent of actin organization. If actin and MTs interact, as suggested previously (Deeks et al. 2010; Wang et al. 2012; Sampathkumar et al. 2011), and if MTs serve as a scaffold for proper actin alignment in response to shape change, then one interpretation of our results could be that in the absence of MTs, actin can more freely explore the space, leading to an apparently less organized network in protoplasts of rectangular shapes. Moreover, because MTs are stiffer filaments, they might have a dominant role on the cytoskeletal organisation in response to cell shape. Nevertheless, because of the absence of cell wall in our system, it is possible that the process by which the actin and MTs usually interact is disturbed.

We have shown that our approach is sufficient to detect defects in the organization of the cytoskeletal network in mutants affecting cytoskeletal dynamics. We found that cell geometry could not rescue the defect of anisotropy in MT network in protoplasts of katanin mutants. It was already shown that a katanin mutant responds less to changes in mechanical stress (Uyttewaal et al. 2012; Sampathkumar, Krupinski, et al. 2014), and that MT reorientation triggered by blue light requires the action of katanin proteins (Lindeboom et al. 2013). In addition, our experiments suggest that a katanin mutant is less responsive to geometrical cues. The previous model of dynamical MTs predicted no influence of cell shape on anisotropy of the network (Mirabet et al. 2018) but experiments suggested the opposite with higher anisotropy for rectangular shapes. By refining the model and adding the severing rule accounting for the katanin action, the simulations recapitulated the effect of shape change on anisotropy of the MTs network. In summary, we propose that in plant protoplasts of *A. thaliana* and in the conditions of our experiments, the MTs self-organize and severing by katanin is required for the response to geometry change. As our 3D simulations of the MT network support our experimental observations, we conclude that geometry is sufficient to explain the specific organization observed.

A recent study highlights the role of CLASP proteins in reorganizing the MT array, notably by stabilizing MTs after severing by katanin (Lindeboom et al. 2019). Future studies could focus on investigating the cytoskeleton organization in our experimental system in protoplasts of CLASP mutants. CLASP proteins have been shown to accumulate at cell edges *in vivo* and suggested to be important for MTs to cross such edges (Ambrose et al. 2011). Whether such CLASP accumulation occurs in protoplasts under geometrical constraints is not clear.

This work is a first step towards assessing quantitatively how cell geometry contributes to the control of cytoskeletal organization in living plant cells. We hope that the approach can be further developed to study how geometry impacts plant cell division or cell polarity.

## Materials and methods

### Seed stock and plant growth conditions

To study cytoskeletal organization, we used green fluorescent protein fused to a MT binding domain (MBD-GFP) (Granger and Cyr 2001), (Marc et al. 1998) and green fluorescent protein fused to an F-actin binding domain (FABD-GFP) (Ketelaar et al. 2004) reporter lines for monitoring the MTs and the actin network, respectively. All *Arabidopsis thaliana* seeds were surface sterilized by ethanol 70%, stratified at 4ºC for 2 days and grown vertically on plates containing Murashige and Skoog medium supplemented with half sucrose under continuous light. For crosses, 10 day old seedlings were transferred into soil and plants were grown under continuous light at 20ºC until flowers appeared, before crossing. For the purpose of this study several crosses were made. *act2-5* plants were crossed with FABD-GFP plants and *bot1-7* plants were crossed with MBD-GFP plants (generated plants were homozygous for the mutant only).

### Callus cell initiation and culture conditions

*Arabidopsis thaliana* (Col-0 accession) calli were prepared from 2 week old seedlings grown *in vitro* under sterile conditions. Roots were collected, chopped into thin sections of ~1 mm in length, and then transferred onto solid callus induction medium (3.8g/L B5 salt mix, 25g/L glucose, 0.625 g/L MES, 1.25mL Gamborg vitamins, pH adjusted to 5.7 with KOH, 62.5 µg/L kinetin, 625 µg/L 2,4D, 10g/L Phytoagar) at 25 °C. The calli were then transferred to a new medium every 2 weeks. Cells from 5 to 9 day old cultures were isolated and used for protoplast generation.

### Protoplast generation

Protoplasts were obtained as previously described (Durand-Smet et al. 2014) by a combination of cell wall degradation and hypo-osmotic shock. Packed cells were gently mixed, in a 15 mL tube, with 5.5 mL of enzyme solution containing 2 mM CaCl2, 2mM MgCl2, 10mM MES, 1 mM L-ascorbic acid, pH 5.5 with KOH, 17 mg/mL Cellulysin (Calbiochem, La Jolla, CA), 17 mg/mL Cellulase RS (Yakult, Co. Ltd., Tokyo, Japan), 0.4 mg/mL Pectolyase Y-23 (Seishin Pharmaceutical Co. Ltd., Nihombashi, Japan), 3.5 mg/mL Bovine Serum Albumin (Sigma, St. Louis, MO), and 600 mOsm with mannitol, sterilized by filtration. Cells were then incubated for 2 hours with rotation shaking (60 rpm) at 25ºC. After 3 min of centrifugation at 800 rpm, the supernatant was discarded and the cells were resuspended in 5mL of washing medium for 5 min (2 mM CaCl2, 2 mM MgCl2, 10 mM MES, pH 5.5 with KOH, 600 mOsm with mannitol). Cells were pelleted again (3 min 800 rpm), the supernatant was removed and 5 mL of hypoosmotic medium (same as washing medium, osmolarity 280 mOsm with mannitol) was added to release protoplasts. After 15 min of gentle shaking (30 rpm), protoplasts were sorted from aggregates by filtration through a 75µm mesh. Fresh protoplasts were transferred into culture medium (PCM, see below) and then directly loaded on the microwell array. We expect that the protoplasts did not regenerate their walls in the time of the experiments (which never exceeded 6 hours after plating except when otherwise specified), as synthesis of a new cell wall generally takes 24 h after cell wall removal (Nagata and Takebe 1970; Cuddihy and Bottino 1982) or 3 days in our system (figS11).

### Protoplast Culture Medium (PCM)

Several culture media have been tested and the one described in (Mathur et al. 1995) had a better yield for protoplast survival in our conditions (see fig S5). We thus decided to use this medium which consisted of: 0.5MS medium containing: IAA 2.0 mg/l 2,4-D 0.5mg/l IPAR 0.5 mg/l and 0.4 M glucose. Fresh protoplasts were then pelleted again (3 min 800 rpm), the supernatant was removed and 1 mL of protoplast culture medium (PCM) was added.

### Drug treatments

For drug treatments, final concentrations of 4µM of Cytochalasin D diluted in 100%DMSO (Sigma, St. Louis, MO) or 20 µM of oryzalin diluted in DMSO (Sigma, St. Louis, MO) were added to protoplasts in PCM for 30min prior to being loaded into the micro-wells (Durand-Smet et al. 2014). Drug-treated protoplasts were observed about one hour after the start of the treatment and imaged in the presence of the drugs. Efficiency of drug treatment is described in the Supplement (fig S7).

### Microwell fabrication

Microchambers were fabricated following standard microfabrication techniques (Weibel et al. 2007). First, shapes of interest were designed with AutoCAD. The CAD file used for this study is available in: https://gitlab.com/slcu/teamHJ/publications/durand_etal_2019

The protoplasts generated with the protocol described here presented diameters of about 25+-8µm on average (standard error). Thus, for each shape, we designed molds with diameters between 15 and 40 µm.

A silicon master with silicon features was created in a clean room following standard micro-lithography techniques. The features were fabricated with a height of 20µm+-0.5µm (figS12). The height of the features was measured with a scanning electron microscope (FEI Sirion). The silicon wafers were then rinsed with ethanol and water and air dried. PCM medium mixed with 1.5% agarose was then poured over the silicon wafer and let cool to gel. Once gelled, the agarose microwells were carefully detached from the wafer.

The microwell array was then deposited on PDMS posts located at the bottom of a plastic petri dish. Fresh PCM was poured such that the air/liquid interface almost reaches the top of the microwells. This was to prevent the agarose from drying throughout the experiment. The protoplasts were then plated on top of the microwells, some protoplasts migrated to the bottom of a microwell by sedimentation (about 1 hour of sedimentation). Protoplasts exhibited shapes dictated by the wells and showed polymerized cytoskeletons several hours following plating, indicating that their general physiology was not perturbed. Only protoplasts that were deformed inside the micro-chambers were taken into account for analysis. Indeed, small protoplasts that fit at the bottom of the chambers without being deformed into square, triangle or elongated shape, remained spherical.

Once the protoplasts were plated into the micro-chambers, a coverslip was carefully added on top of the array before imaging under an upright microscope for observation (fig1). The protoplasts are usually observed 30 min to 1h after being plated, thus the organization we observed forms on a time scale faster than hours. Moreover, the specific organization observed here in case of the rectangular shape is persistent in time and could still be observed 24h after the protoplasts were plated (figS13). In our experiments we thus monitor the steady state organization of the cytoskeletal network.

### Confocal image acquisition and image analysis

Protoplasts lodged in micro-chambers were imaged with a Zeiss 780 or Zeiss 880 Airyscan confocal laser scanning microscope with a 63X oil objective and Z-stacks of cells with 0.18 μm intervals were obtained for 3D reconstruction.

Images taken with the 780 confocal were processed with ImageJ (Schindelin et al. 2012), http://reb.info.nih.gov/ij).

Zen 2.3 software was used to process the images acquired with the 880 Airyscan confocal microscope. The software processes all Airy channels in order to obtain images with enhanced spatial resolution in 3D (Korobchevskaya et al. 2017).

After processing, z-stacks were projected on one plane with maximum intensity projection with FIJI to create 2D images.

The FIJI plugin Fibriltool (Boudaoud et al. 2014) was used in order to quantify the average orientation and the anisotropy of the network. The average orientation gives information on the direction of the network, while the anisotropy gives an estimation on how well the filaments are aligned with each other.

Region of interest were drawn manually to define the contours of the cells on the 2D pictures and nematic tensors of MT and actin filament arrays inside the region of interest (ROI) were obtained using FibrilTool. Fibriltool directly provides the average orientation of the network inside the ROI and the anisotropy of the network with a score between 0 and 1 for every cell. The anisotropy is given by *q* = *n*_*1*_− *n*_*2*_ where *n*_*1*_ and *n*_*2*_ are the eigenvalues of the nematic tensor **n** with **n** = **t** ⊗ **t**. The unit vector 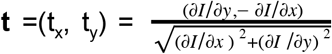 is the tangent to the putative fibrillar structures and *I*(x,y) is the pixel intensity level in the picture.

Further information on the validation of the quantification extracted from the pictures is provided in Supplement (figS14).

### Statistical analysis

For anisotropy comparison, two sample Kolmogorov–Smirnov (KS) tests were performed. The KS tests used are the ones currently in the Python Scipy library (E. Jones et al. 2001).

Kuiper tests were run on angular circular data. To that end, uniform distributions of the same number of angles as in the compared dataset were generated and a two sample Kuiper test was run (code available at https://github.com/aarchiba/kuiper/blob/master/kuiper.py). Results of KS tests on the same data sets are provided for information.

Results of tests with p-values higher than 0.05 were considered not significant.

### Numerical simulation description and parameter choice

Our numerical model is an adaptation of that developed in Mirabet et. al 2018 (Mirabet et al. 2018), which we will outline briefly here. Each tubulin contained within a MT is modeled as a constant length vector, with MTs as multi-segment vectors. The plus-end of a MT is marked to grow, shrink or remain static and the minus-end to shrink or remain static. Shrinking remove successive unit vectors representing tubulin from the corresponding MT end. Growth adds a unit vector to the corresponding end, with the direction matching that of the previous tubulin modified by a small amount of randomness. MTs are contained within a defined domain (an ellipsoid, square, cube and cuboid were considered in Mirabet et al 2018). MTs are either strongly anchored where they are restricted to the membrane surface after first contact with the membrane; or weakly anchored where MTs are prevented from crossing the membrane by redirecting their growth direction along the membrane (before the addition of the random component) but can still reenter the cell interior after contact with the membrane, and additionally when they are within 10nm of the domain surface, if the angle between the surface and MT is bigger than α_*anch*_ the MT starts shrinking. MTs were nucleated 80nm within and parallel to the cell surface at a constant rate, *n*_*p*_, per time step per unit surface. The MT minus-end detaches with probability, *n*_*s*_, per nucleation site per time step, at which stage the MT starts shrinking from the minus-end causing treadmilling or shrinkage depending on plus-end behaviour. For MT interactions Mirabet et al imposed that when a growing MTs is within distance *d* of the next closest MT: if the angle between them is less than α it bundles with the direction of the new unit vector tubulin aligning with the neighbouring MT, but if the angle is greater than α an induced catastrophe occurs and the growing end changed to shrinking.

Rather than always causing an induced catastrophe when the MTs interact at angles greater than α, we introduced a probability (1 − *P*_*cat*_) that the MTs continues to grow but will be cut at the crossover at a later time with probability *P*_*cross*_ per tubulin per time step. When a MT plus end undergoes a growth event, we first determine if its nearest neighbour MT is within distance *d*_1_. If it is, and the angle is below the threshold α the MT bundles with its growth direction changing to match that of the existing MT (following Mirabet et al 2018). If the angle is larger than the threshold, we assume this MT will be crossing the neighbouring MT. This causes an induced catastrophe with probability *P*_*cat*_, in which case the MT growing plus-end changes to shrinking. Otherwise, the MT keeps growing and this element is added to a cut list, and the nearest-neighbor tubulin element that we assume is the one crossed is also recorded. At each growth step, every element in the cut list is severed with a probability *P*_*cross*_ but only if its nearest-neighbour tubulin element still exists (this ensures the MT will not be severed if the MT it crossed over has been destroyed). Severing breaks the MT into two at the cut element, with both MTs shrinking from their cut ends (figS4 (Anon n.d.)). Given that the model is fully 3D and MT adjacency is defined by a distance threshold, this method allows adjacent tubulin on the same MT to be recorded as crossing for the same cross-over (so tested for induced catastrophes or added to the cut list). Thus, relatively small probabilities can correspond to relatively large levels of cutting or induced catastrophes, for example, if *P*_*cat*_ = 0.001 then assuming a MT is checked for an induced catastrophe at 10 tubulin during a cross-over then the probability of an induced catastrophe at that MT crossover is 1 − (1 − 0.001)^10^ ≈ 0.01, nearly ten times larger. (We also tested the alternative to computationally impose that only the first tubulin in a continuous line of tubulin classed as a crossover is marked for an induced catastrophe or for cutting, leading to a decrease in the number of elements in the cut-list and a shallower anisotropy trend with increasing cut probability, but no change to the conclusion within this paper. This condition is not imposed for any of the simulations presented in this paper.)

Our Simulations were run for *n* = 10000 or *n* = 3000 steps. With a simulation time step of approximately 0.1 to 0.2 seconds this corresponds to a real time of approximately 15-30min and 5-10 minutes respectively. The number of repeats were *N* = 50 or *N* = 20. We used the pre-existing square and rectangle domains outlined in Mirabet et. al. 2018. We assume zero probability of MT plus-end growth randomly changing to shrinkage per timestep per MT, *P*_*s*_ = 0. We assume weak anchoring and take α = 0.7. At the start of the simulation *n*_*init*_ = 30 MTs are nucleated, with subsequent nucleations occuring at a rate of nucleations per time step corresponding to 2.5-5 nucleations per second (or *n*_*p*_ ≈ 1).

To compare to experimental anisotropy results calculated from single-plane images, we calculate anisotropy in 2D, on two (parallel) opposite faces of our domain. The rectangle and square both align with the coordinate axes, so to restrict to these two planes, we calculate the maximum and minimum tubulin position in the out-of-plane coordinate axis and include tubulin with an out-of-plane coordinate within *d*_*aniso*_ =1.36µm (corresponding to 170 computational units) of either value. Tubulin directional vectors are projected onto the plane of the face to reduce them to 2D. Our anisotropy measure is the *S*_2_ (Deinum et al. 2011) where

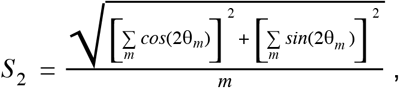

where *m* is the total number of tubulin in all the MTs and θ_*m*_ the angle between the tubulin and a reference direction (for every tubulin the same length), where 0≤*S*_2_≤1 and *S*_2_ = 0 corresponds to isotropy and *S*_2_ = 1 to perfect alignment. The orientation of alignment (Deinum et al. 2011) is calculated as

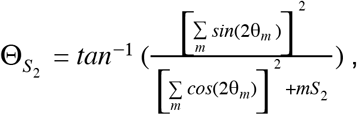

where 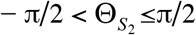.

The MT simulations were implemented in c++ with anisotropy calculations performed in Python on the output vtk files.

### data availability

Original confocal data is available via the University of Cambridge Research Data Repository (doi:...), code used for the analysis and 3D MT simulations are available on the Sainsbury Laboratory gitlab repository (https:///gitlab.com/slcu/teamhj/publications/durand_etal_2019, https://gitlab.com/slcu/teamhj/Tubulaton).

## Acknowledgments

We thank all members from E. M. Meyerowitz and H. Jönsson groups for their advice during the course of the project. We thank Sarah Robinson and Olivier Hamant for helpful discussions. We thank Arun Sampathkumar and Francois Nedelec for providing suggestions on the manuscript.

PDS was funded by global fellowship Marie Sklodowska-Curie (PlantCellMech H2020), TS and HJ Gatsby (GAT3395/PR4, PRXXX/YY) and HFSPO (grant RGP0009/2018). The work in the Meyerowitz laboratory at the California Institute of Technology was partially funded by the Howard Hughes Medical Institute.

## supplementary

**Figure S1:**
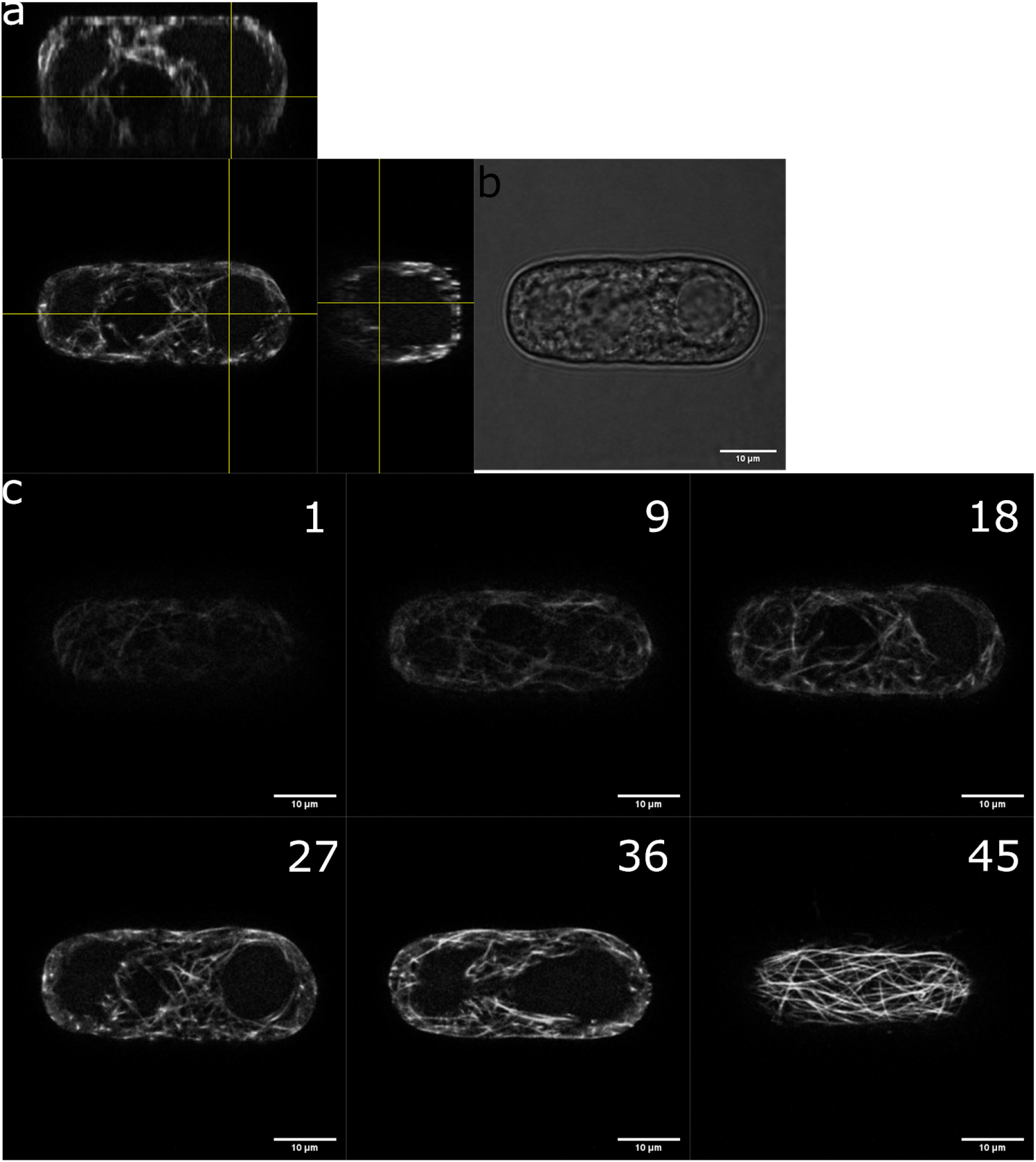
a, orthogonal views of a protoplast expressing MBD-GFP reporter in a rectangular micro-well. b, bright field picture of the same protoplast. c, different focus planes of the same protoplast.

**Figure S2:**
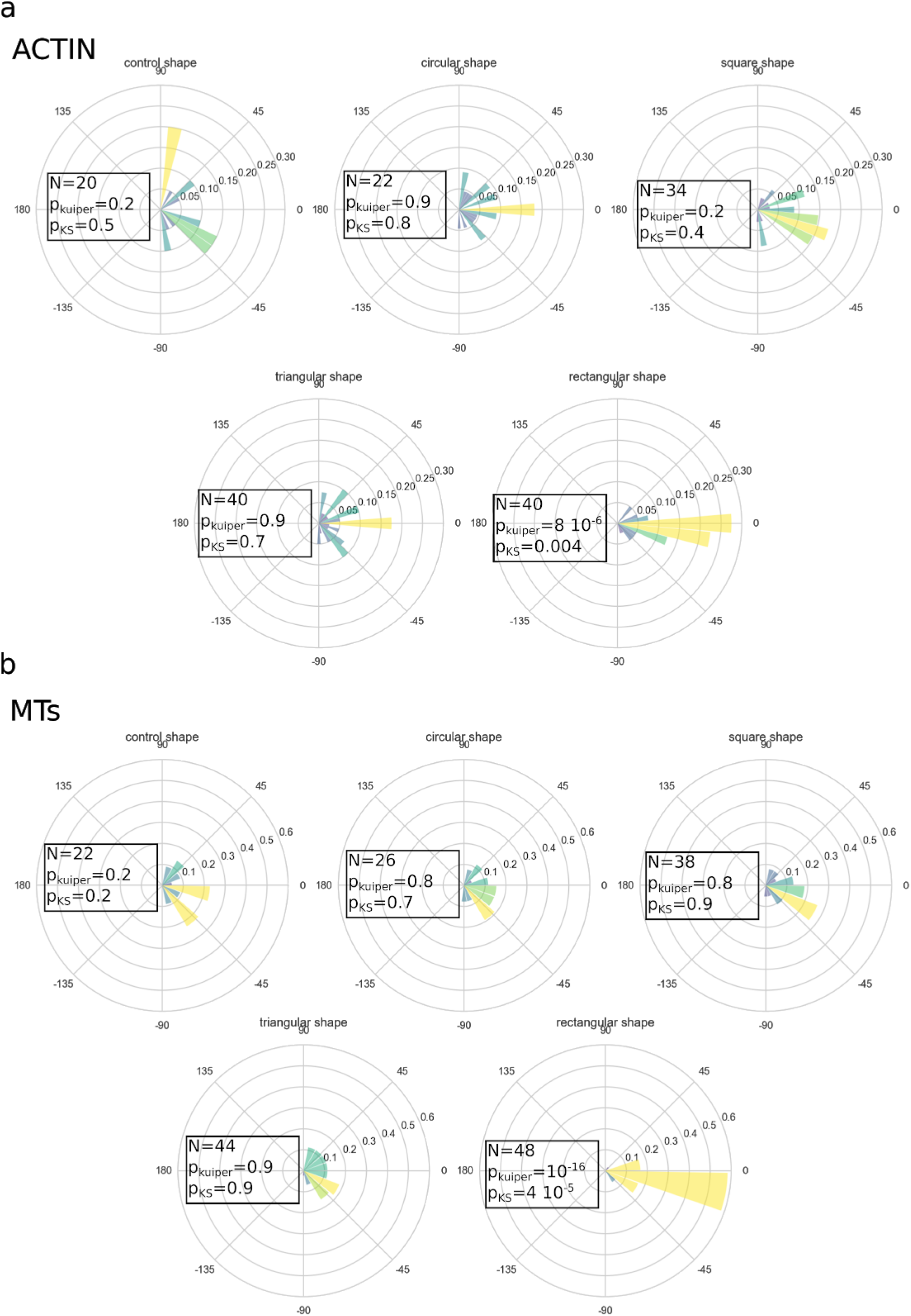
(a) Angular histograms of the average angle of the actin network in protoplasts in different geometries. (b) Angular histograms of the average angle of the microtubule network in protoplasts in different geometries. Pkuiper and pKS are the pvalues returned by Kuiper test and Kolmogorov-Smirnov test.

**Figure S3:**
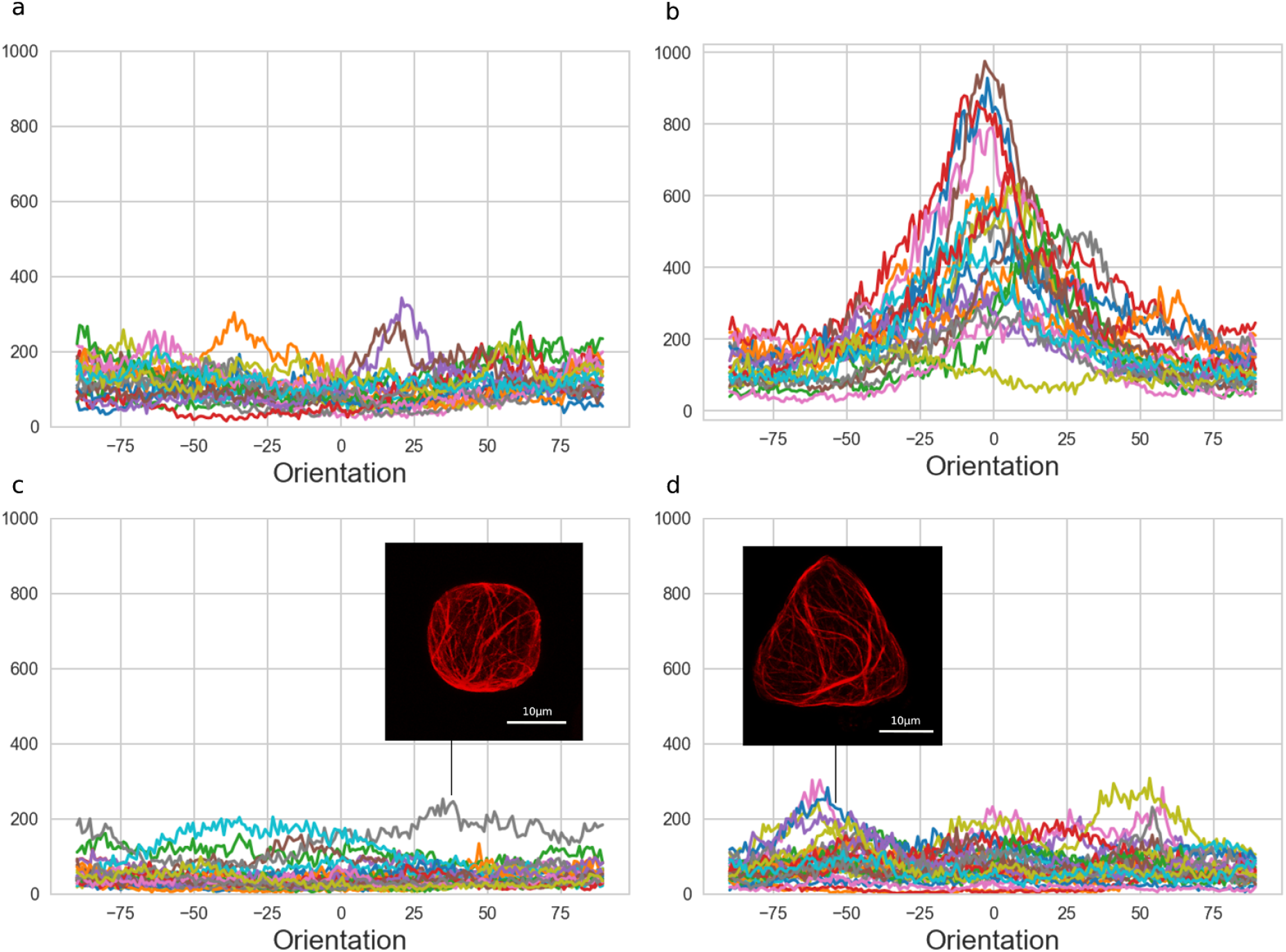
Angle distributions of microtubules of all cells in (a) control shapes (b) rectangular shapes (c) cubic shapes and (d) triangular shapes. The distributions were generated with OrientationJ, a Fiji plugin (http://bigwww.epfl.ch/demo/orientation/)(Rezakhaniha et al. 2012) a local window size of 2 pixels and a cubic spline gradient were chosen to run the analysis for the orientation of the fibers. (c) The insert picture shows the microtubule network of a protoplast in a cubic micro-well where the microtubules were organized along the diagonals (6/38 protoplasts presented such an organization). (d) The insert picture shows the microtubule network of a protoplast in a triangular micro-well where the microtubules were organized along the edges (5/30 protoplasts presented such an organization).

**Figure S4:**
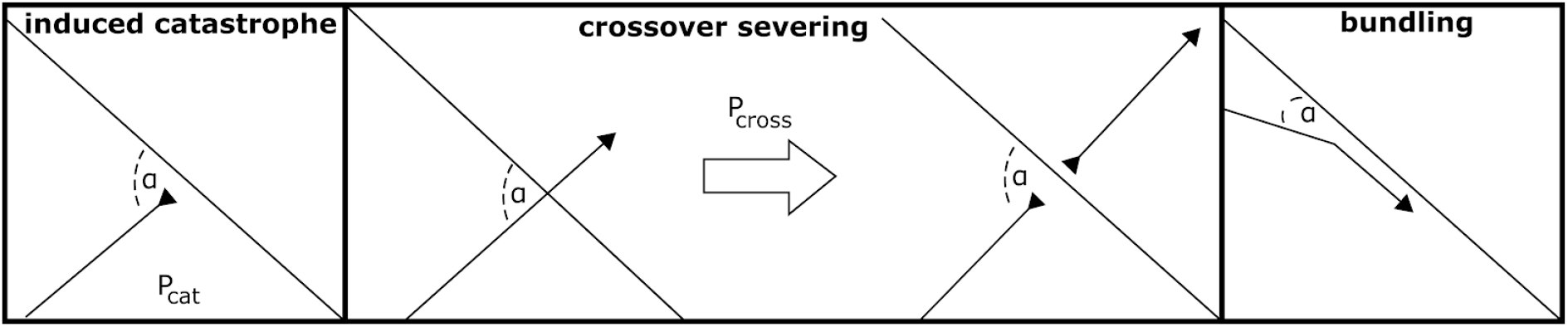
Diagram showing the different types of microtubule interactions in the numerical simulations.

**Figure S5:**
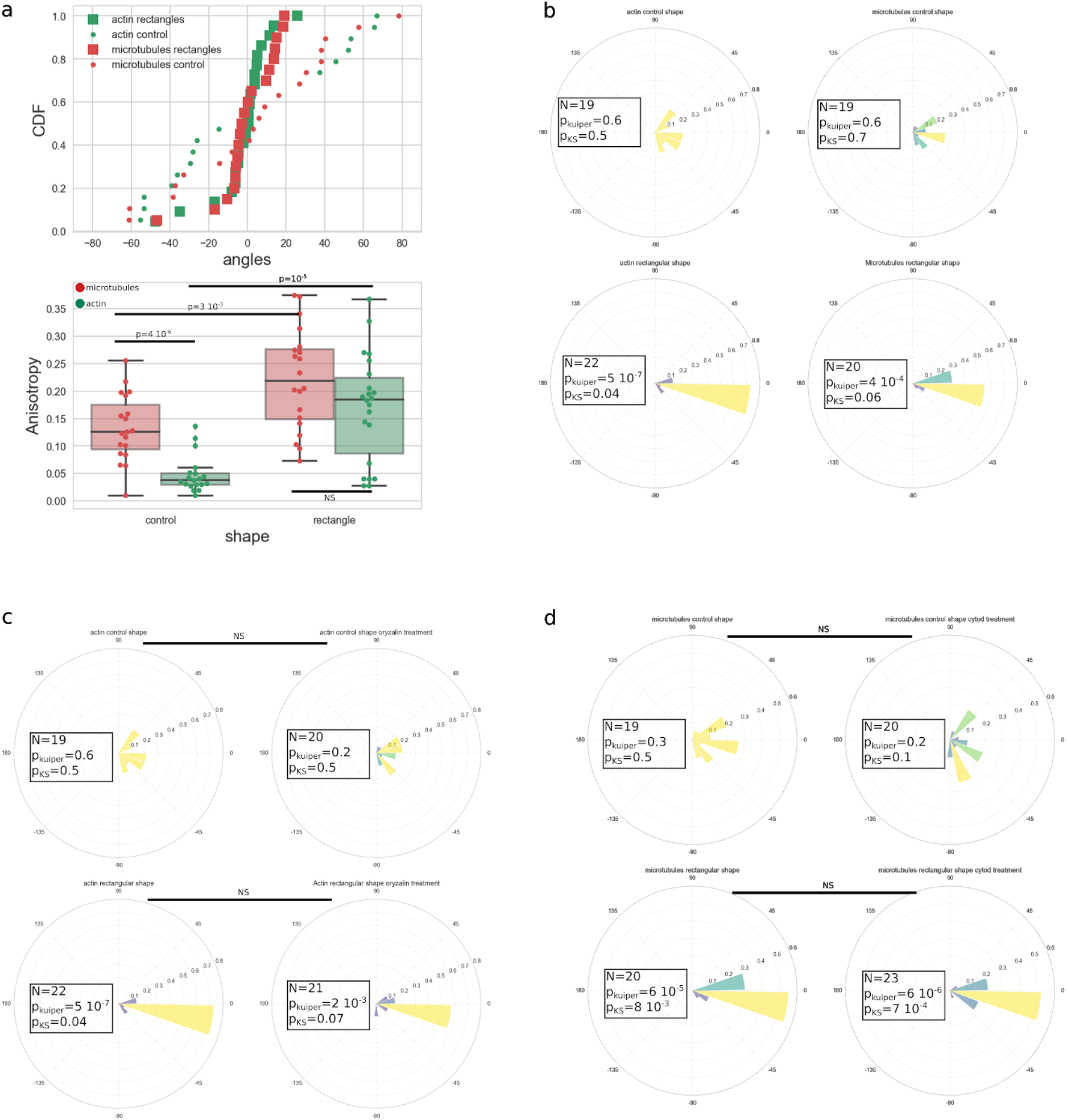
Actin and microtubule networks organized similarly in protoplasts confined in elongated geometry. (a) Actin and microtubule networks are organized similarly in protoplasts confined in elongated geometry. Cumulative distributions of the average angle of the actin (green) and microtubule (red) networks in rectangular microwells (square markers) and in control shape (spherical protoplasts. circle markers). Box plot comparing the anisotropy of the microtubule and actin networks in rectangular and control shapes. The actin network is always less ordered that the microtubule network but in both cases, the networks are significantly more ordered in rectangular shapes than in spheres. (b) Angular histogram of the average angle of the microtubule (right) and actin (left) networks in protoplasts in control geometry (top) and in rectangular micro wells (bottom). Both actin and microtubules organize along the main axis of the rectangular micro-wells. (c) Angular histograms of the average angle of the actin network in protoplasts in control geometry (top) and in rectangular micro wells (bottom) for untreated protoplasts (right) and protoplasts treated with 20μM of the microtubule depolymerization agent oryzalin (left). (d) Angular histograms of the average angle of the microtubule network in protoplasts in control geometry (top) and in rectangular micro wells (bottom) for untreated protoplasts (right) and protoplasts treated with 4μM of the actin depolymerization agent CytochalasinD (left).

**Figure S6:**
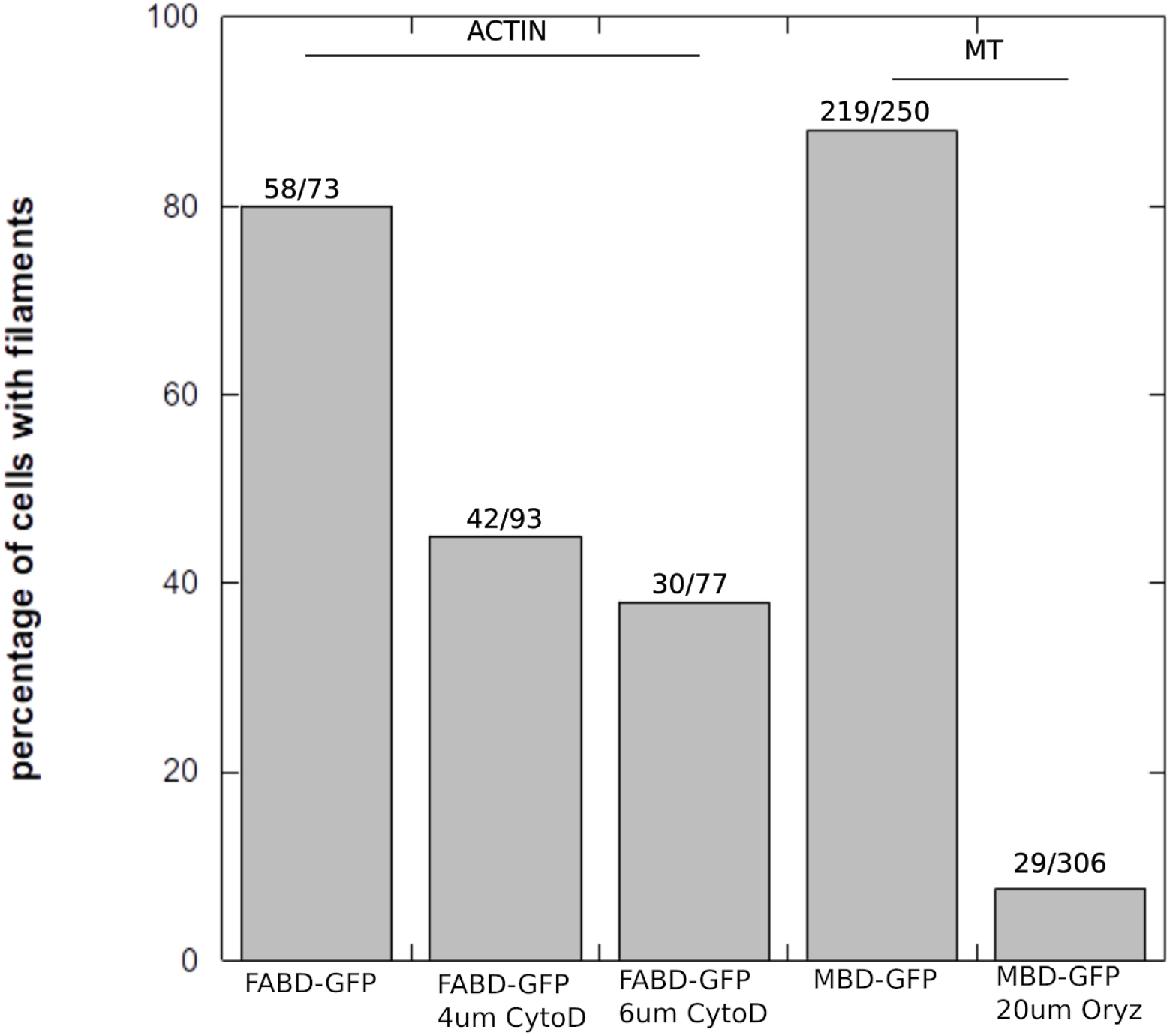
Number of protoplasts with apparent filaments after drug treatments. Protoplasts were generated, then treated with drugs for 30 min as described in the protocol in Materials and Methods, and then protoplasts were mounted on a hemocytometer for observation using a confocal microscope in order to check the presence of actin filaments (in FABD-GFP lines) or microtubules (in MBD-GFP lines).

**Figure S8:**
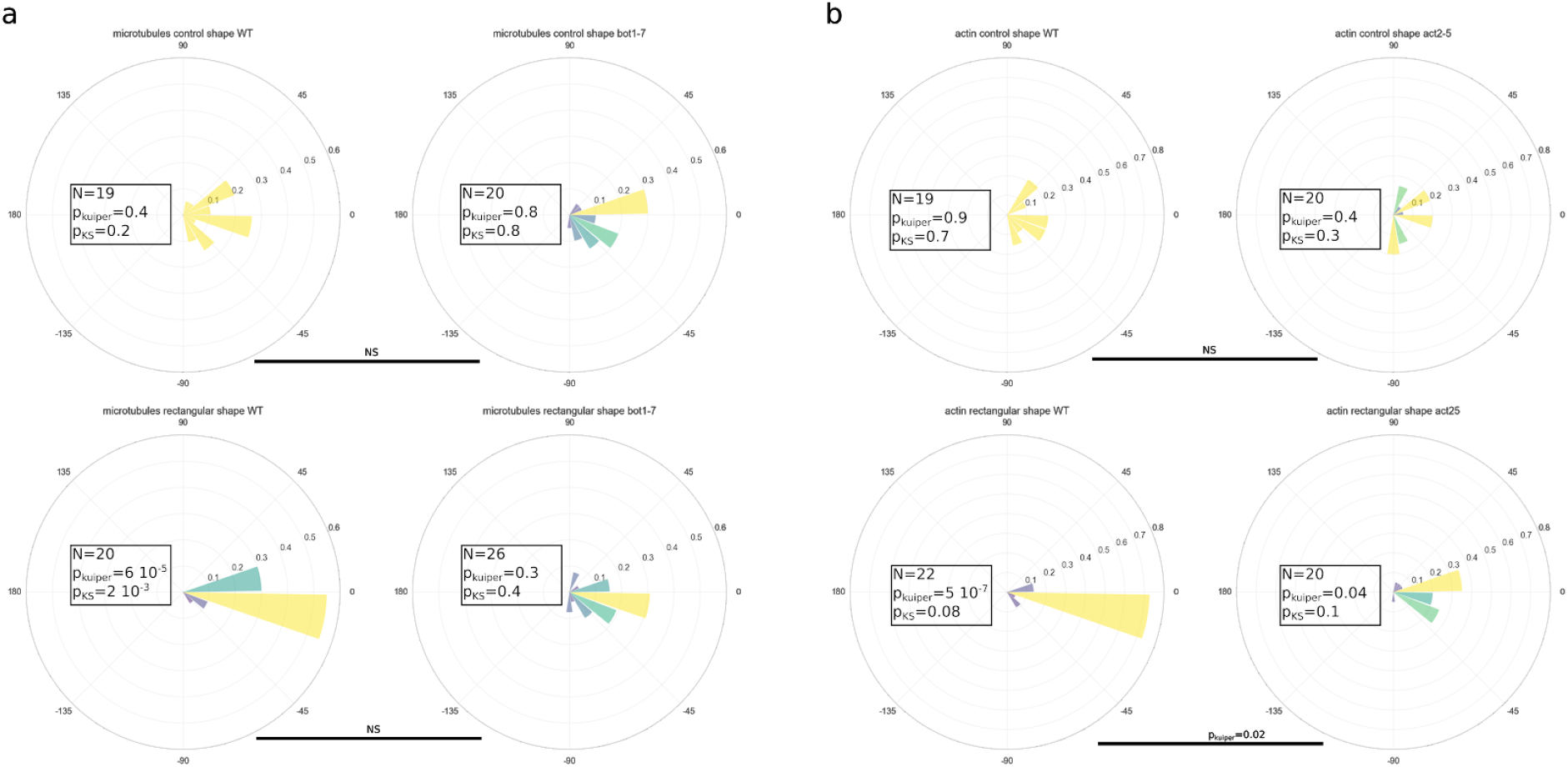
(a) Angular histograms of the average angle of the microtubule network in protoplasts in control geometry (top) and in rectangular micro wells (bottom) for wild type protoplasts (right) and protoplasts in the *bot1-7* background mutant (left). (b) Angular histograms of the average angle of the actin network in protoplasts in control geometry (top) and in rectangular micro wells (bottom) for wild type protoplasts (right) and protoplasts in the *act2-5* background mutant (left).

**Figure S9:**
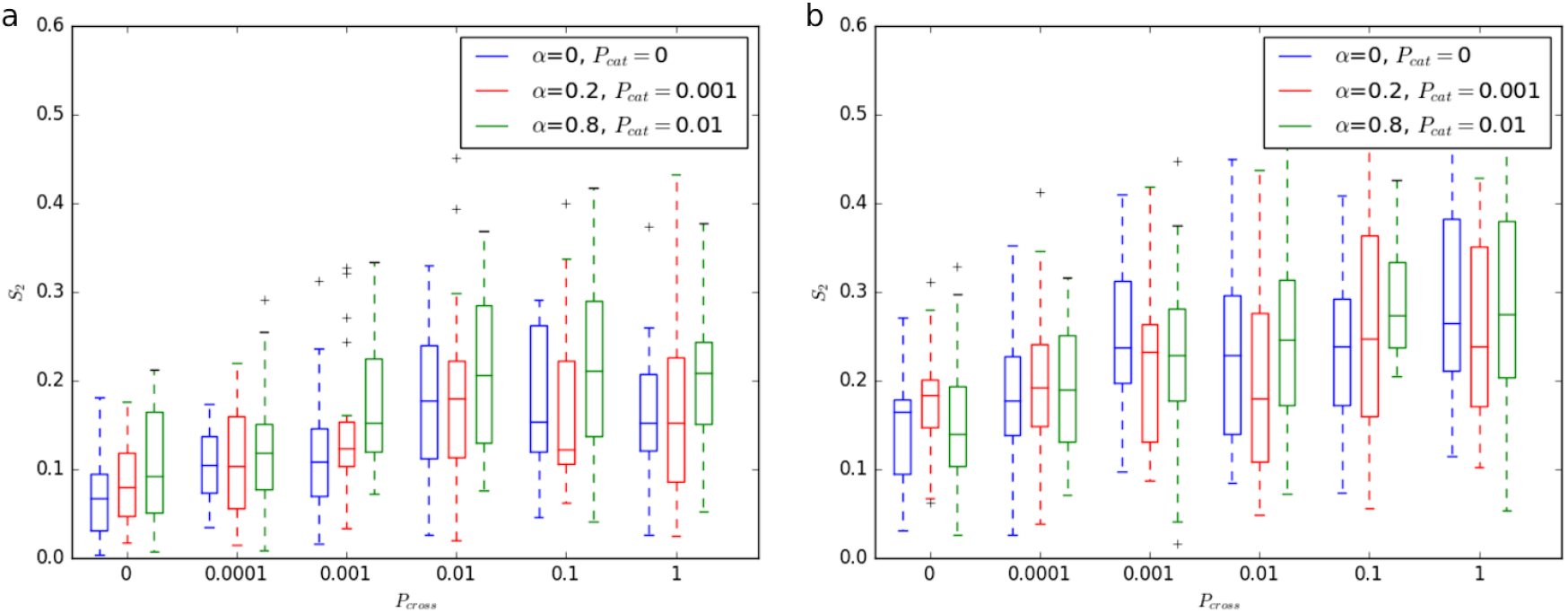
Computational simulations comparing anisotropy for increasing *P*_*cross*_ for three cases with different probabilities of induced catastrophe and bundling angle for square (a) and rectangular (b) domains.Three cases for different *P*_*cat*_ and α are shown. We first considered the case with no induced catastrophes or bundling. The anisotropy is lower at smaller values of *P*_*cross*_. We then considered two more realistic cases with both bundling and induced catastrophes included, where a similar trend is observed. However, at very large *P*_*cat*_, varying *P*_*cross*_ has little or no effect on anisotropy.

**Figure S10:**
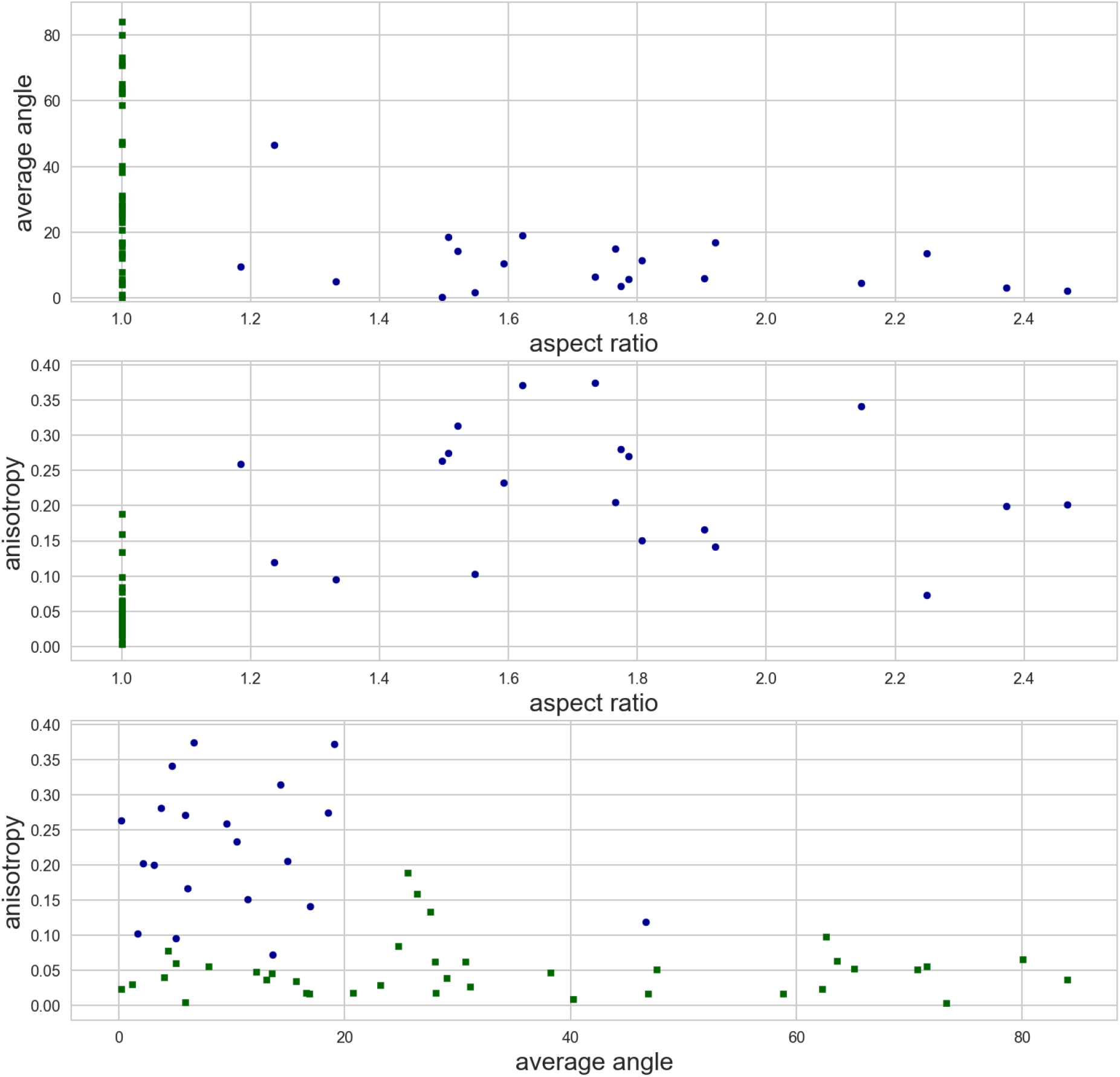
Evolution of the average angle (upper graph) and anisotropy (middle graph) of the MT network as a function of the aspect ratio of the protoplasts (aspect ratio of 1 corresponds to protoplasts confined in square shapes). Lower graph: anisotropy and average angle of the MT network in protoplasts confined in rectangular shape (circle markers) and square shape (square markers).

**Figure S11:**
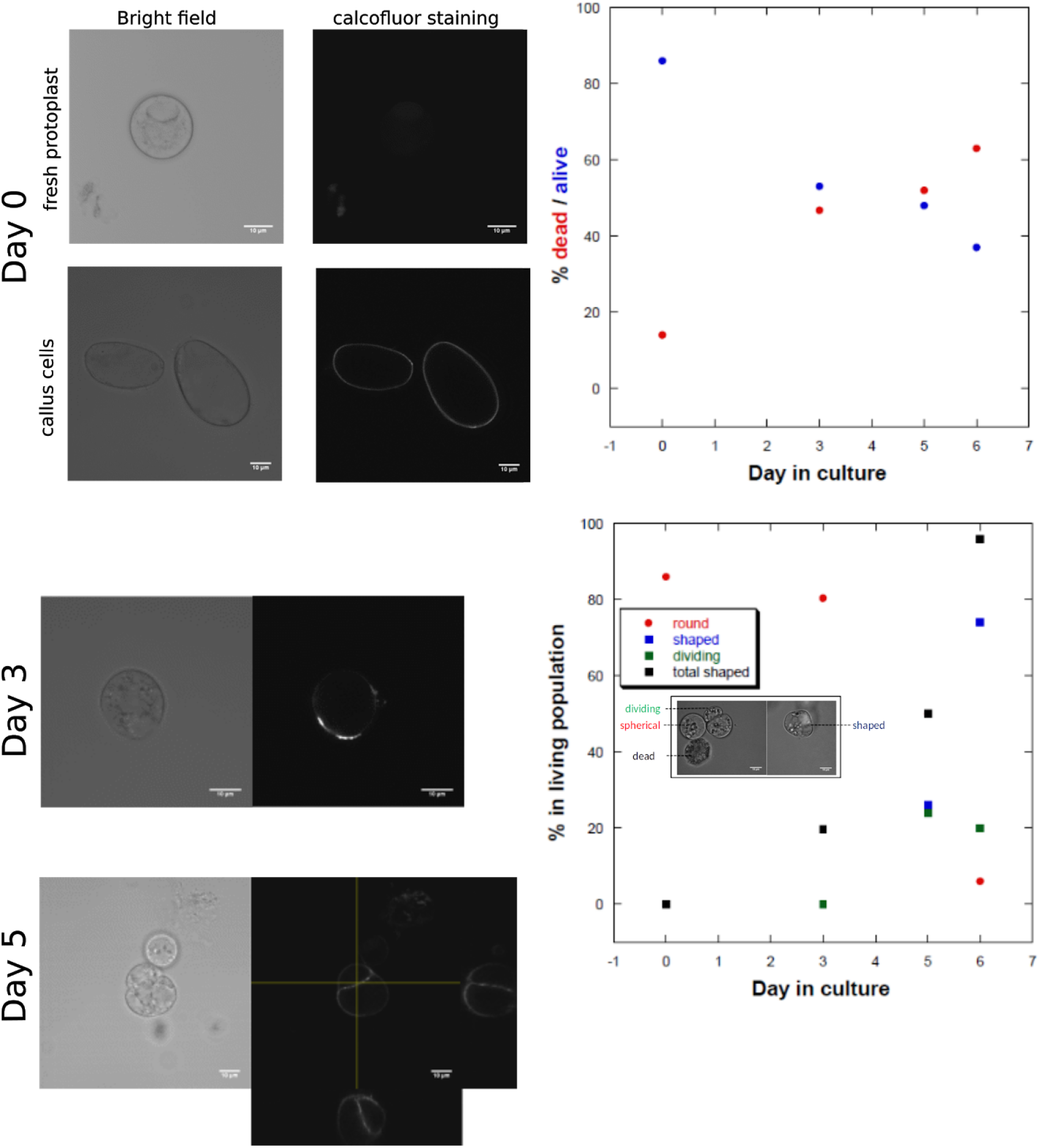
left: calcofluor cell wall staining (0.01% CW) in fresh protoplast and callus cells. Fresh protoplasts do not exhibit cellulose fluorescence, in contrast to callus cells where accumulation of cellulose can be observed in the conditions of staining. After 3 days and 5 days in culture, new cell walls start appearing in the protoplast culture, and some dividing cells can be observed, showing that the medium chosen allows protoplasts to survive for several days. The graphics on the right show the percentage of living protoplasts in the culture as a function of days in culture. While the living protoplast population decreases and reaches 40%, the number of protoplasts with new cell walls (shaped) or dividing in that population increases (bottom panel). For this regeneration and survival assay, protoplasts were cultivated in 5cm Petri dishes, plated at a density of 1.5 x10^6^ cells/ml in the culture medium described in Materials and Methods. Counting of living/dead and shaped/dividing/round cells was performed with a hemocytometer.

**Figure S12:**
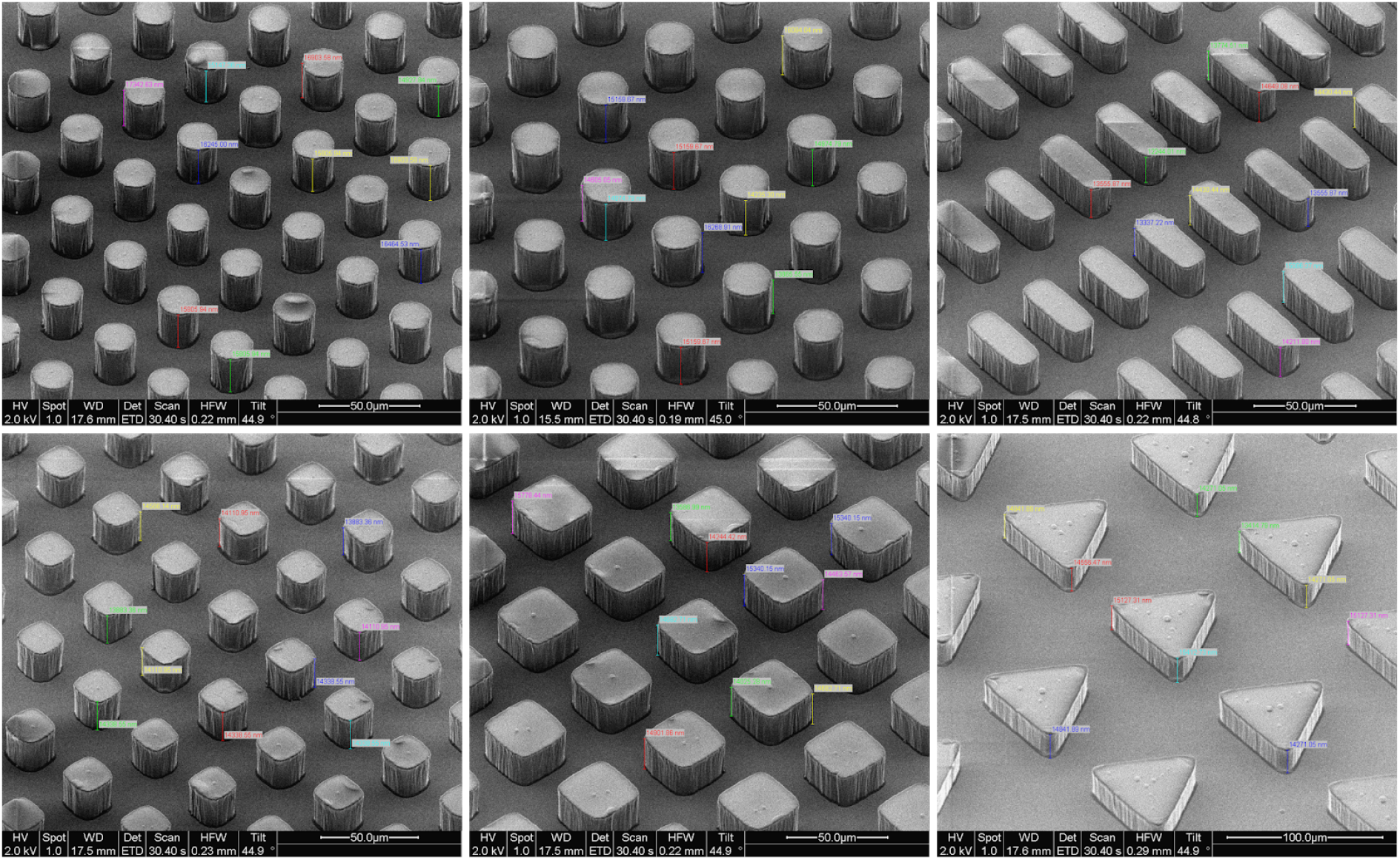
Scanning Electron Microscopy pictures of silicon masters, with SU8 photoresist features, used as molds for micro-chamber fabrication. The features are made of SU8-2025, an epoxy-based photoresist designed for micromachining (MicroChem) and the real average height is 20+-0.3um.(real height= height/sinα with α the tilt angle, 44 degree on the above pictures).

**Figure S13:**
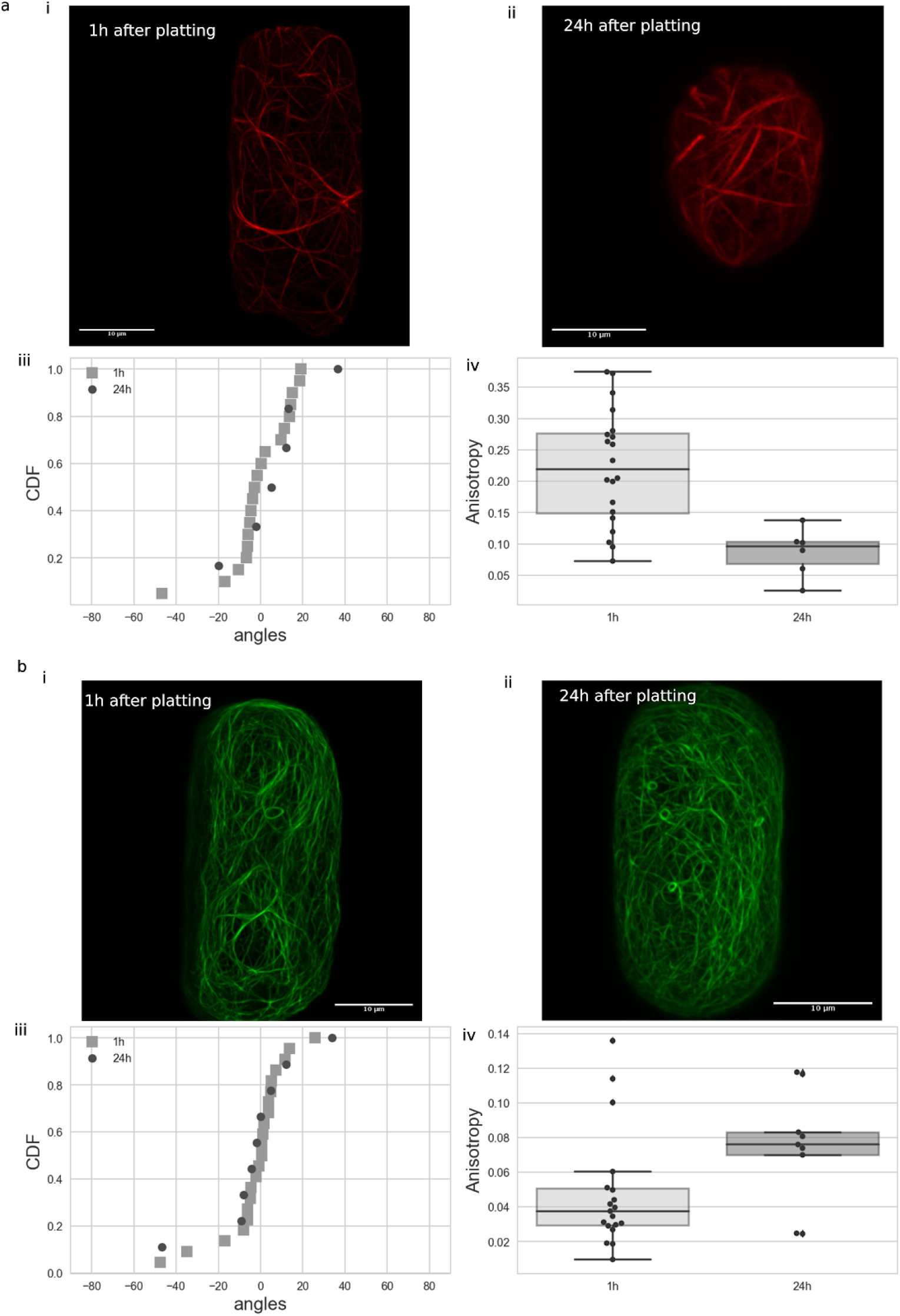
Quantification of the organisation of the microtubule (a) network and actin (b) network in protoplasts confined in rectangular shapes at 1h or 24h after plating.

**Figure S14:**
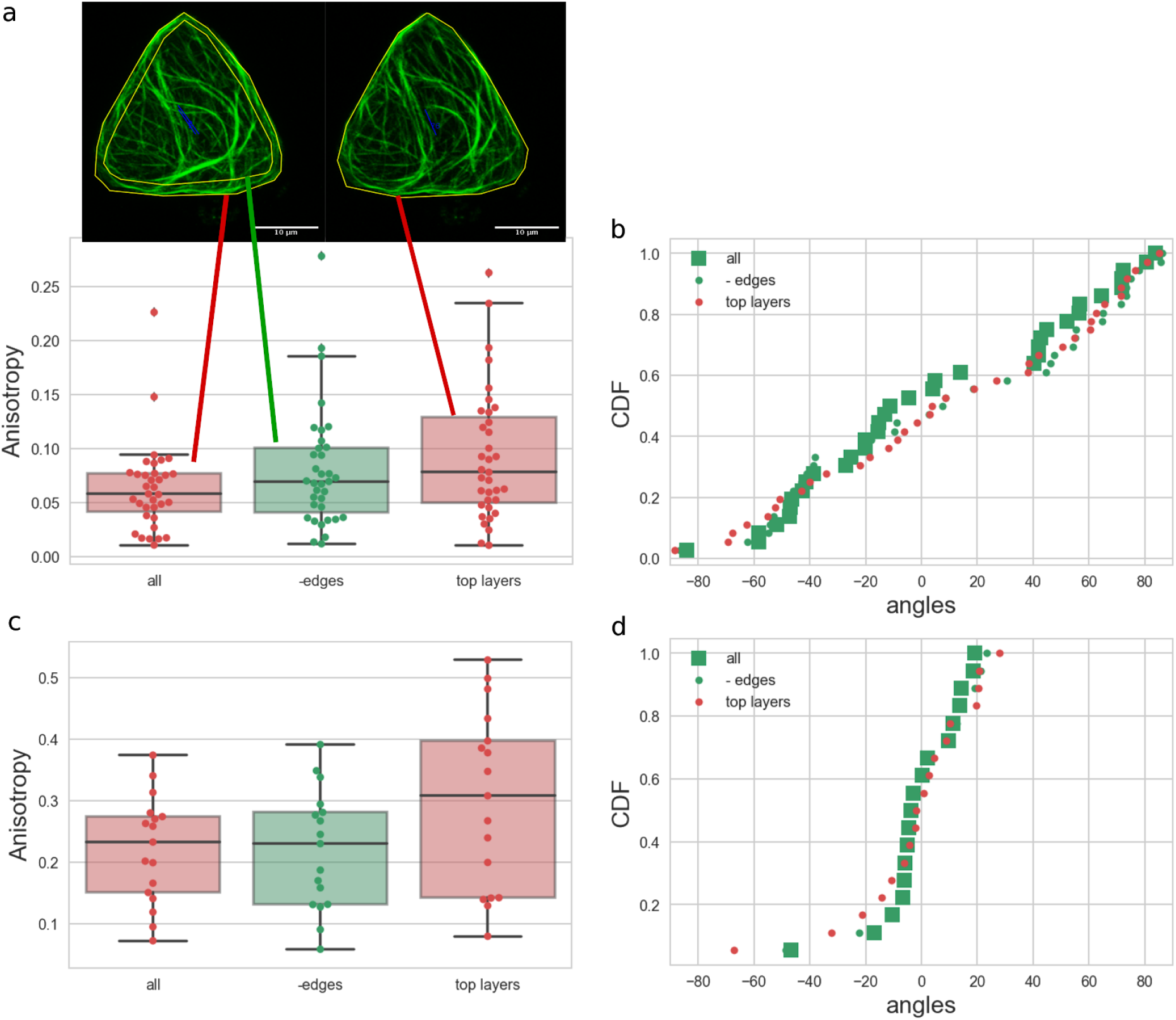
Comparison of the quantification of the microtubule network organization in protoplasts in triangular shapes(a-b) and rectangular shapes(c-d). (a-c)anisotropy distributions obtained from Fibriltool analysis on whole pictures (all focus plans from z-stack projected onto one plane), pictures with the border of the cells removed (to see the effect of accumulation at the border of the shape due to the maximum intensity projection) and when only the 10 top focus planes from the z-stacks were taken for the maximum intensity projection. (b-d)distribution of the average angles obtained from Fibriltool analysis on whole pictures, pictures without borders, or pictures with top layers only. for all cases, the distributions between the 3 types of analysis (all, -edges or top layers) did not present significant differences when statistical tests were performed.

**Table S1:**
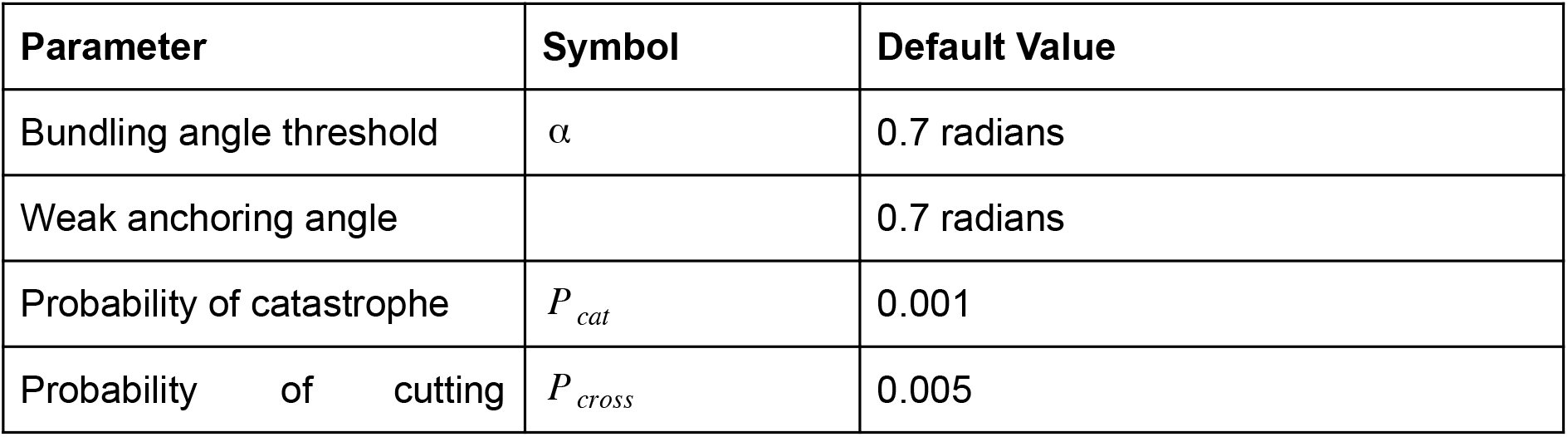
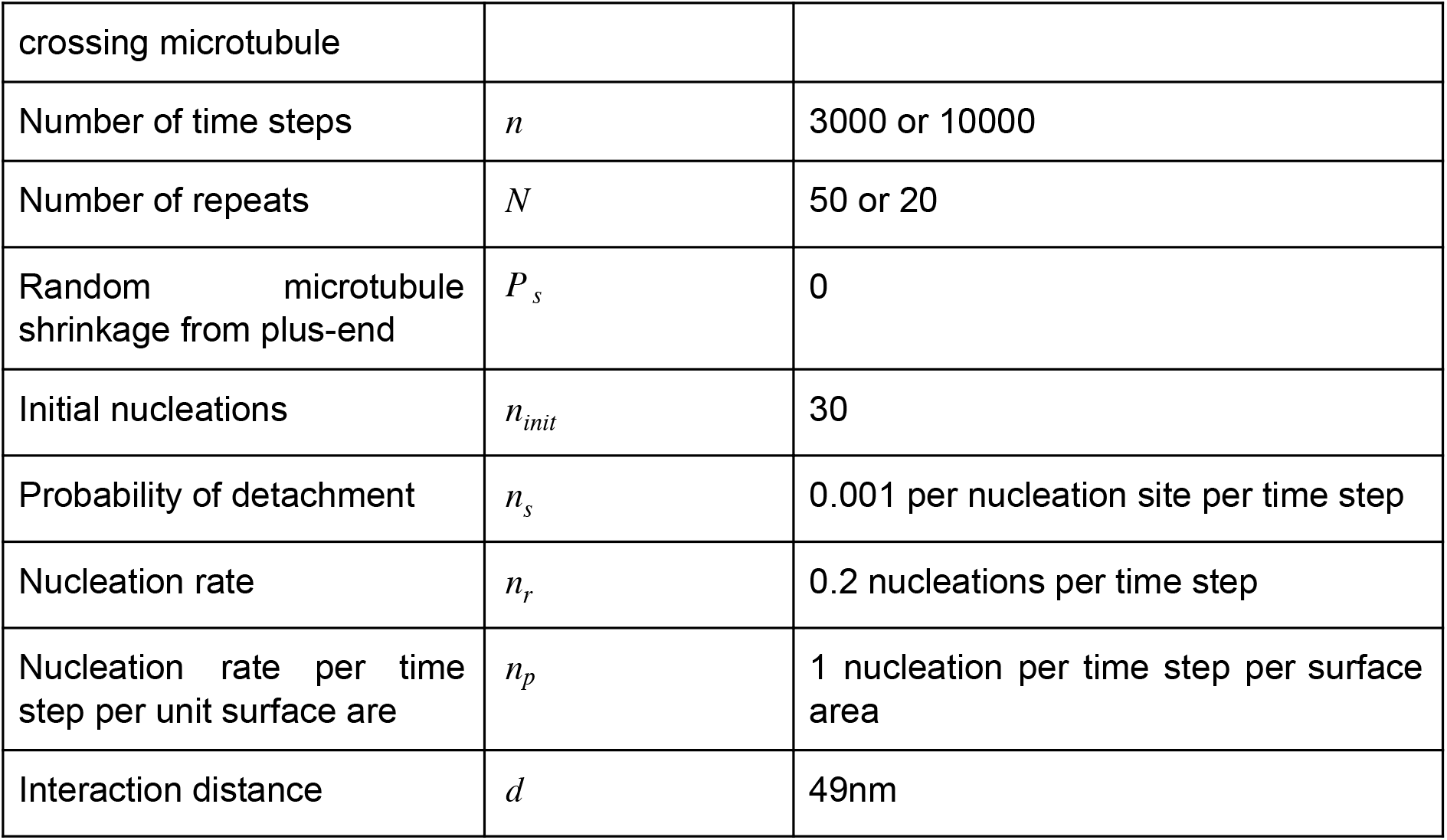
Computational parameters used in the numerical simulations

